# Lipid monolayers adsorbing mRNA as models for mRNA enclosed in lipid nanoparticles for transfection

**DOI:** 10.1101/2025.08.01.667014

**Authors:** Miriam Grava, Konrad Weber, Joshua Reed, Benjamin Weber, Chen Shen, Heinrich Haas, Emanuel Schneck

**Affiliations:** Institute for Condensed Matter Physics, TU Darmstadt, Hochschulstraße 8, 64289 Darmstadt, Germany; BioNTech SE, An der Goldgrube 12, Mainz 55131, Germany; Deutsches Elektronen-Synchrotron DESY, Notkestrasse 85, 22607 Hamburg, Germany; Department of Biopharmaceutics and Pharmaceutical Technology, Johannes Gutenberg University, 55128 Mainz, Germany

**Keywords:** LNP, nanoparticles, mRNA, drug delivery, Langmuir monolayers, x-ray scattering, x-ray fluorescence

## Abstract

In lipid-based mRNA pharmaceuticals applicable for vaccination, cancer therapy, and other types of therapeutics, the negatively charged RNA is embedded in nanoparticles containing positively charged or ionizable lipids. The local cohesion at the lipid-RNA interface plays an important role for the stability and biological activity of the resulting nanoparticles. Ions and other buffer components in the RNA’s immediate surroundings may affect RNA stability against hydrolysis and its binding strength to the oppositely charged lipid layers, which is relevant for release after cellular uptake and endosomal processing. Here, we use Langmuir monolayers at the air/water interface as well-defined experimental models to study the local molecular organization of mRNA adsorbed to lipid layers. Binding of mRNA in the presence of monovalent and divalent ions from the aqueous phase to monolayers consisting of cationic transfection lipids and zwitterionic phospholipids is investigated with synchrotron-based x-ray fluorescence and scattering techniques. The experiments provide detailed insights into the structure of the adsorbed layers as well as the fractions of all molecular moieties contributing to the interfacial electrostatic balance, specifically phosphates from RNA, phosphates from phospholipids in the monolayer, and elemental counterions. The mRNA forms discrete electron-dense layers tightly bound to the lipid membrane, where cationic ions accumulate at the interface together with the mRNA. This leads to an excess of anionic nucleotides relative to cationic lipids at the interface, which is more pronounced in the presence of divalent compared to monovalent cations. The quantitative information on local interfacial moieties obtained here may constitute a valuable basis for the evaluation of quality aspects of nanoparticles comprising mRNA in the bulk phase.

## Introduction

Validated by the success of the mRNA-based COVID-19 vaccines, pharmaceutical products based on mRNA have been gaining great scientific and public interest.^1^ In addition to infectious disease vaccines, potential applications include cancer therapy, protein replacement, and vaccination against bacterial infections.^2,3^ To exploit the full potential of mRNA-based pharmaceutical products, aspects related to quality and structure-function correlation of nanoparticulate mRNA pharmaceutical products need to be better understood.^4–6^ In particular, product characteristics which are a consequence of their colloidal nature, and the processes involved in the self-assembly of different molecular components into entities with supramolecular organization have been underrepresented in the commonly applied panel of control assays. For extended characerization, application of quantitative biophysical methods has reached a certain level of general acceptance. Size distribution profiles and size-dependent structural parameters can be obtained by various methods, including nanotracking analysis (NTA), size exclusion chromatography (SEC), asymmetrical flow field flow fractionation (AF4), analytical ultracentrifugation (AUC), Taylor dispersion analysis (TDA), tunable resistive pulse sensing (TRPS) and others. ^6–8^ Electron microscopy, X-ray scattering, and neutron scattering are frequently employed for the investigation of structure and morphology at molecular-scale resolution.^6,9,10^ However, for a broader acceptance of these methods, standardized protocols for data analysis and interpretation are required.

In the present study, we focus on the local conditions directly at the mRNA-lipid interface. In lipid nanoparticles (Fig. 1 A) the mRNA is present in the hydrophilic region between two adjacent lipid layers, comprising lipid head groups, water, buffer and excipient molecules. ^11^ Electrostatic interactions between the negatively charged phosphate groups of the mRNA and the positively charged lipid head groups are essential for maintaining the supramolecular organization in the nanoparticles. The conditions in this compartment are substantially different from those in the bulk phase, for example with regard to molecular composition, mobility, and pH, which has tremendous influence on chemical stability and on processes around endosomal trafficking, and therefore on biological activity.^12–14^ Chemical degradation processes, such as lipid and RNA hydrolysis or adduct formation, which are key limiting factors for product shelf life, sensitively depend on the exact localization of molecular moieties with respect to each other. ^14^ Since it is difficult to obtain such high resolution structural information using bulk dispersion samples, we have chosen Langmuir lipid monolayers at the air/water interface^15^ as well-defined model systems to study these phenomena with the help of synchrotron-based grazing-incidence X-ray scattering techniques (Fig. 1 B) that have previously been used successfully to determine the pH-dependent protonation degree of ionizable lipids^16^ and to study the interaction of DNA to lipid layers.^17,18^ Grazing-incidence X-ray off-specular scattering (GIXOS) ^19–21^ yields the electron density profiles perpendicular to the interface and thus resolves the interfacial layer structure.^22^ The simultaneous use of total-reflection X-ray fluorescence (TRXF)^23,24^ spectroscopy allows quantifying the surface densities of chemical elements (here: of phosphorous atoms in lipid and RNA and of the counterions). In the present work, by spreading mixtures of transfection lipids and phospholipids on a pharmaceutically relevant subphase containing mRNA, we studied the binding of mRNA and ions to the lipid layer surface. We determined the amount of mRNA relative to the fraction of transfection lipids in the monolayer membrane, the density of ions and their effect on mRNA binding and conformation, and revealed how these moieties contribute to the electrostatic balance at the interface.

**Figure 1:**
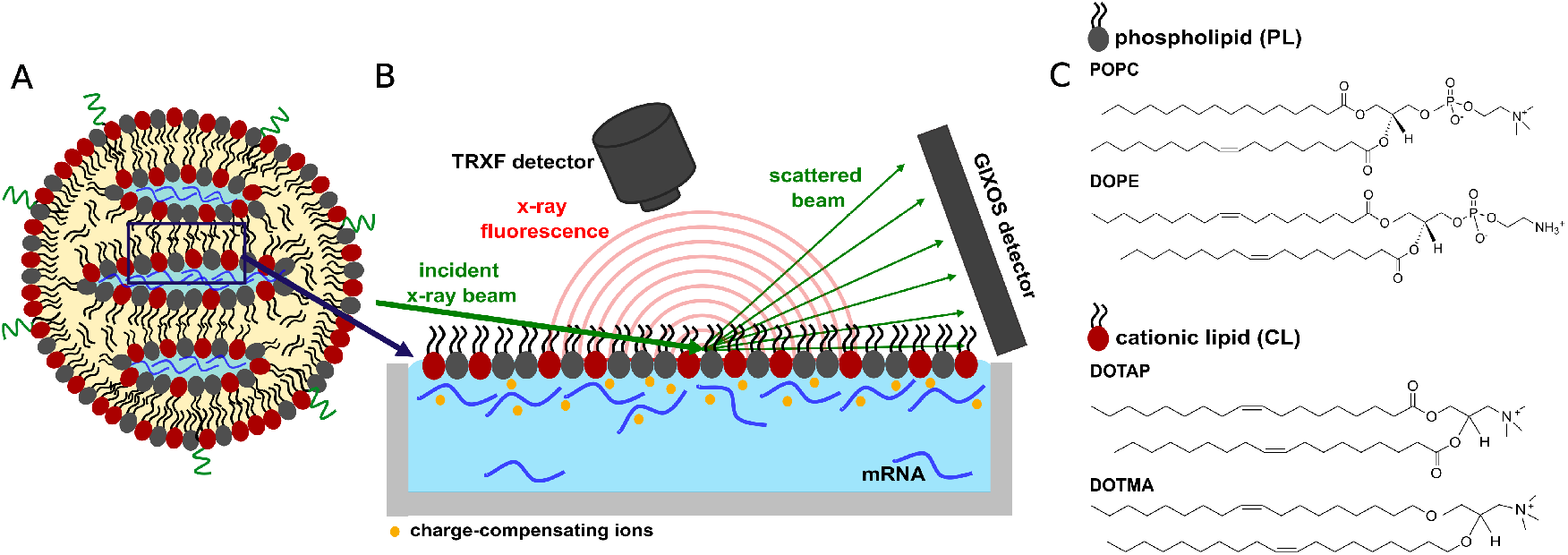
(A) Schematic illustration of a lipid-based nanoparticle featuring lamellar lipid structures loaded with mRNA. (B) Schematic illustration of the experimental setup consisting of a lipid monolayer at the air/water interface, where ions and mRNA are dissolved in the subphase. The interface is illuminated with a synchrotron X-ray beam at grazing incidence under total-reflection conditions. The electron density profiles and the elemental/molecular compositions of the interfacial layers are determined from simultaneous measurements of GIXOS and TRXF, respectively. (C) Molecular structures of the phospholipids POPC and DOPE and of the transfection lipids DOTAP and DOTMA. The monolayers are 1:1 or 2:1 mixtures (mol/mol) of transfection lipids and phospholipids (see main text).

## Results

Fig. 1 B schematically illustrates the experimental setup used for our investigations. The lipid-mRNA interface inside the lipid nanoparticles is mimicked by a planar lipid monolayer (or “half a lipid membrane”) floating at the air-water interface. Mixtures of zwitterionic phospholipids (PL) and a cationic lipids (CL) are used to represent the lipid composition of the nanoparticles. The setup allows to vary the average area per lipid, *A*_lip_, and to monitor the associated change of the lateral pressure Π. Ions and mRNA are dissolved in the aqueous subphase from where they can adsorb to the lipid layer. The adsorption layer is studied with high accuracy through simultaneous application of two grazing-incidence X-ray scattering techniques, GIXOS and TRXF. Their surface sensitivity is based on the total external reflection of the incident X-ray beam that thus only illuminates the immediate vicinity of the air/water interface (see methods section).

GIXOS measures the electron density profile perpendicular to the interface and is therefore sensitive to changes of the electron density profile due to the adsorption of RNA to the lipid layer. TRXF is sensitive to the interfacial elemental composition. It yields the interfacial concentrations of phosphorus (P) atoms belonging to the PL (from which the lateral density of the CL can be derived indirectly) and of mRNA, being a polynucleotide with each nucleotide comprising one P atom. Moreover, TRXF yields the concentrations of anions and cations accumulating in the interfacial region.

For the lipid monolayers, we used mixtures of PL and CL, similar to those used in formulations, which have proven efficacy for delivery of tumor antigens to dendritic cells for cancer immunotherapy.^3,25–28^ Their chemical structures are shown in Fig. 1 C and the full chemical names are found in the Methods section. We investigated a 1:1 mixture of the CL DOTAP with the PL POPC, termed “DOTAP/POPC” in the following, a 1:1 mixture of DOTAP with the PL DOPE, termed “DOTAP/DOPE”, and a 2:1 mixture of the CL DOTMA with DOPE, termed “DOTMA/DOPE”.

Lipid monolayers at the air/water interface were formed by spreading the lipid mixtures dissolved in chloroform on the surface of an aqueous subphase. The subphase contained either only monovalent ions (5 mM KCl or KBr), or it additionally contained mRNA at a concentration of 0.1 mg/mL (corresponding to a nucleotide concentration of about 0.3 mM). In a further set of experiments, which we refer to as “with Ca^2+^” in the following, in addition to the monovalent ions and the mRNA, the subphase contained also divalent cations (1 mM CaCl_2_ or CaBr_2_). After spreading and equilibration, the monolayers were laterally compressed to a surface pressure of Π = 30 mN/m, where the molecular packing is known to represent that in tension-free lipid bilayers well.^29^ *A*_lip_ at this lateral pressure can be estimated from the pressure–area isotherms, as shown in the Supporting Information (Fig. S1). However, in view of several potential error sources in context with the monolayer preparation, in this study we determine *A*_lip_ from the lateral density of P atoms with the help of TRXF (see further below), which we consider more reliable. As shown in the Supporting Information (Table S1), injection experiments demonstrated that *A*_lip_ at Π = 30 mN/m is virtually unaffected by the interaction with polynucleotides.

### Electron density profiles as obtained with GIXOS

Fig. 2 A shows a representative set of GIXOS curves, obtained with a DOTMA/DOPE monolayer on three different subphases, one containing only monovalent ions, one containing monovalent ions and mRNA, and one containing monovalent and ions and mRNA as well as divalent calcium ions. Experiments with the two other lipid mixtures (DOTAP/POPC and DOTAP/DOPE) gave qualitatively similar results, which are presented in the Supporting Information (Figs. S2 and S3). The pronounced intensity oscillations along the *q*_*z*_-axis indicate the presence of layers with defined thicknesses, homogeneous electron densities, and defined boundaries at the air/water interface (see Methods section). The *q*_*z*_-position of the first intensity minimum (indicated with arrows in the figure) is reciprocally correlated with the total layer thickness. The significant shift to lower *q*_*z*_ in the presence of mRNA reflects an increase in the layer thickness, which can be attributed to the adsorbed mRNA layer. The further shift in the presence of Ca^2+^ indicates a significantly higher thickness of the adsorbed mRNA layer than in the presence of only monovalent ions.

**Figure 2:**
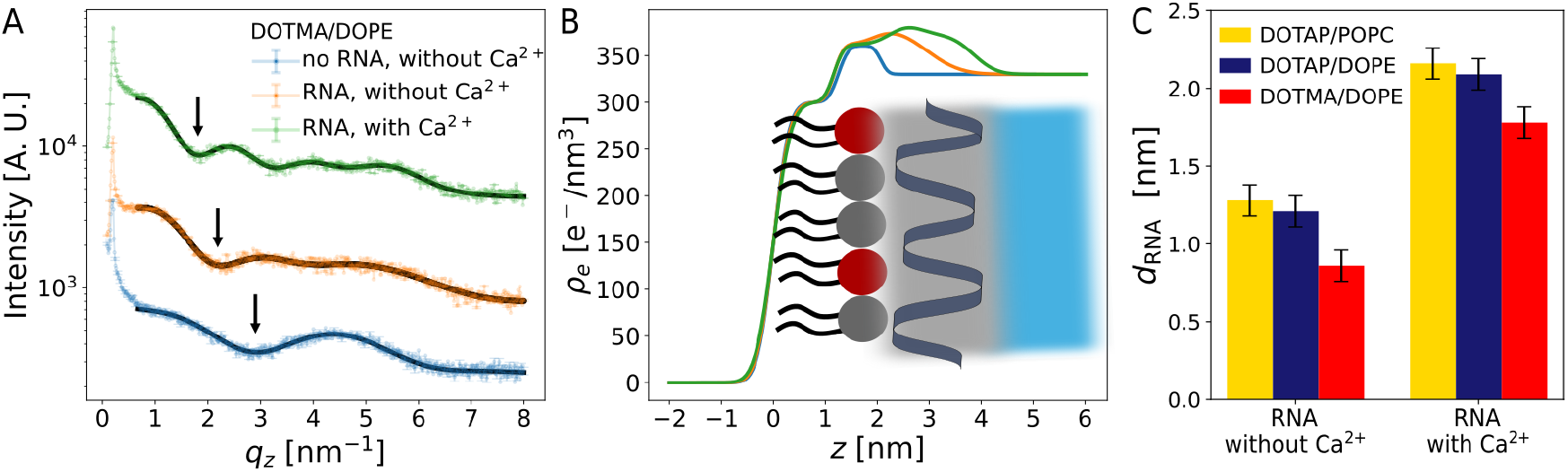
(A) Experimental (symbols) and modeled (black solid lines) GIXOS data for a DOTMA/DOPE monolayer on different subphases. (B) Corresponding electron density profiles *ρ*(*z*). For clarity, the curves in panel (A) are vertically offset in the semi-logarithmic plot through multiplication with suitable factors (5 and 20). Arrows indicate the position of the first minimum, which reflects the overall layer thickness. Data for DOTAP/POPC and DOTAP/DOPE are presented in the Supporting Information (Figs. S2 and S3). (C) RNA adsorption layer thicknesses *d*_RNA_ deduced from the GIXOS data for all lipid compositions in the presence and absence of calcium. See Methods section for the definition of the error bars.

For quantitative analysis of the GIXOS data, the electron density profiles perpendicular to the interface were modeled as sets of stratified layers of adjustable thickness values *d* and electron density values *ρ*. The electron density gradient between the layers was described with error functions with roughness parameters *ε* (corresponding to one standard deviation). The associated theoretical GIXOS curves, modeled from the electron density profiles as described in the Methods section, are shown in Fig. 2 A as solid lines superimposed to the data points, where one can see that the experimental data are well reproduced. In agreement with earlier studies ^16,24,30^ the lipid monolayer in the absence of mRNA (Fig. 2 A, bottom) can be conveniently described with two layers, one for the electron-poor hydrocarbon chains (HC) and one for the electron-rich headgroups (HG). The obtained best-matching layer thicknesses *d*_HC_ and *d*_HG_ are presented in Table 1, and the complete set of layer parameters is given in the Supporting Information (Tables S2-S4).

**Table 1:**
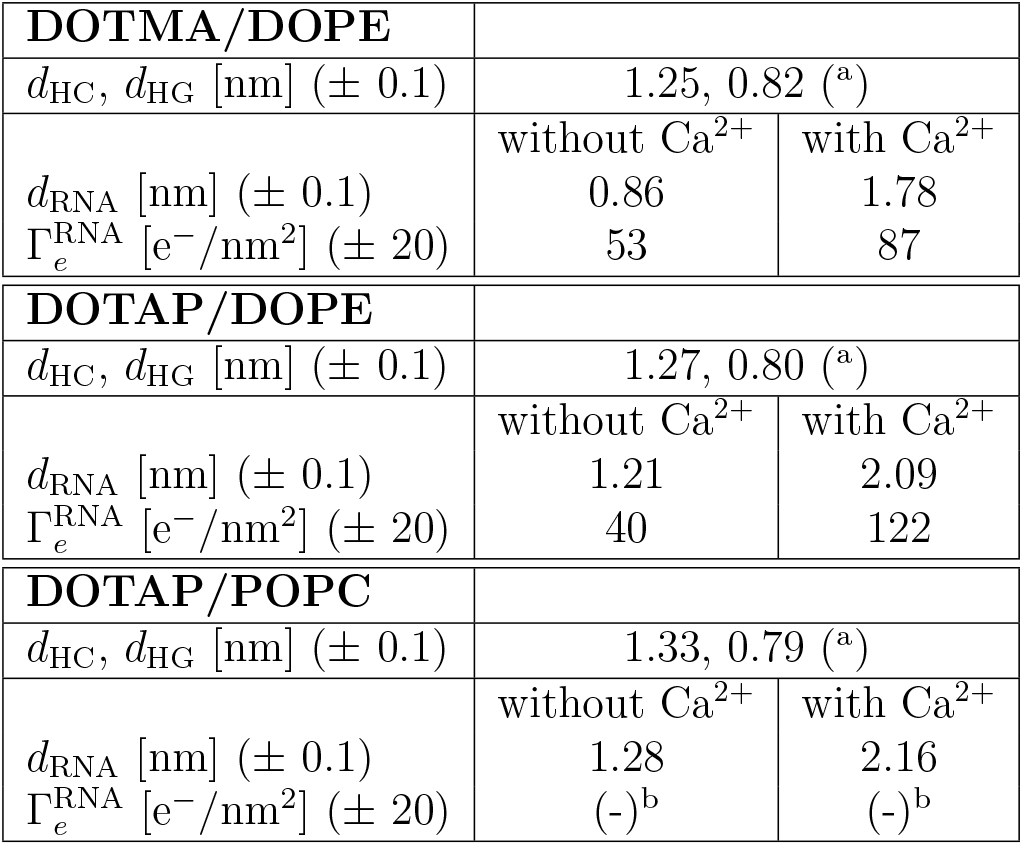
Monolayer characteristics in terms of the hydrocarbon and head group layer thicknesses *d*_HC_ and *d*_HG_ as well as RNA adsorption layer thicknesses *d*_RNA_ and RNA-associated excess electron densities 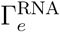, all deduced from the GIXOS data. ^a^Refers to the monolayer in the absence of mRNA. ^b^Not quantified when Br^−^ is the anion (see main text). See Methods section for the definition of the error estimates.

GIXOS data obtained in the presence of mRNA (shown for the DOTMA/DOPE monolayer in Fig. 2 A, middle and top) were modeled with two additional layers (adjacent to the head groups) with different electron density to account for the adsorbed mRNA layer. A single layer for the RNA with uniform electron density was typically found insufficient to reproduce the experimental GIXOS data with the model, supposedly due to different packing of the mRNA moieties in direct contact with the lipid membrane compared to those facing to the subphase. In the following, we focus on the overall changes of the electron density profile upon mRNA binding. We calculate the overall mRNA layer thickness *d*_RNA_, given by the difference between total layer thickness with and without mRNA, and the amount of RNA-related excess electrons per unit area, 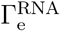, obtained by integrating the electron density difference along the profiles in the presence and absence of mRNA (for details see Methods section). 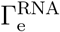 not only comprises the electrons from the mRNA, but also contributions from the charge-compensating ions redistributed upon mRNA adsorption. These effects have to be taken into account for the interpretation of 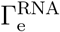, in particular when different monovalent salts (Br^−^ vs. Cl^−^) are present. For better comparability we therefore report 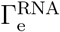 only for the two lipid compositions where KCl was used as monovalent salt. The obtained values of *d*_RNA_ and 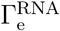 are summarized in Table 1, and *d*_RNA_ is shown as a comparative plot in Fig. 2 C. Consistently for all lipid layer compositions, in the presence of only monovalent ions, *d*_RNA_ is of the order of 1 nm, while in the presence of divalent cations it is about twice as thick (*≈* 2 nm). Consistently, the electron excess 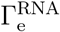 due to the electron-rich mRNA is significantly larger in the presence of Ca^2+^ than in its absence. The experimentally observed electron density of the mRNA layer is given by the contributions from all components, namely the mRNA itself, the hydrating water molecules, and the ions that participate in the overall charge balance. A quantitative discussion of the electron densities will therefore require information about the layers’ molecular composition, which can be obtained by TRXF (see following section).

Summarizing the GIXOS results, as graphically represented in Fig. 2, we found the mRNA binding to lipid layers comprising transfection lipids in the form of discrete layers with sharp interfaces, which underlines the tight binding to the oppositely charged surface. In this sense, the structure resembles the organization of entities with defined size, like proteins, ^31^ bound to monolayers, rather than polymers which may have more random coil organization towards the subphase.

### Composition of the adsorbed mRNA layer as obtained by TRXF

Fig. 3 shows a representative set of TRXF spectra, obtained with a DOTMA/DOPE monolayer on a subphase with monovalent ions only, and with such a monolayer on a subphase with mRNA, in the presence of monovalent ions and, additionally, divalent cations (calcium). The dominant X-ray fluorescence emission lines of the chemical elements of interest (here: P, Cl, K, and Ca) are indicated. The spectra obtained with the other lipid compositions are similar and shown in the Supporting Information (Fig. S4 and S5), together with a representation spectrum in the full energy range (Fig. S6). For samples involving Br^−^ as the anion, the emission line of Br instead of Cl is observed. Details of the TRXF measurements and analysis are presented in the Methods section.

**Figure 3:**
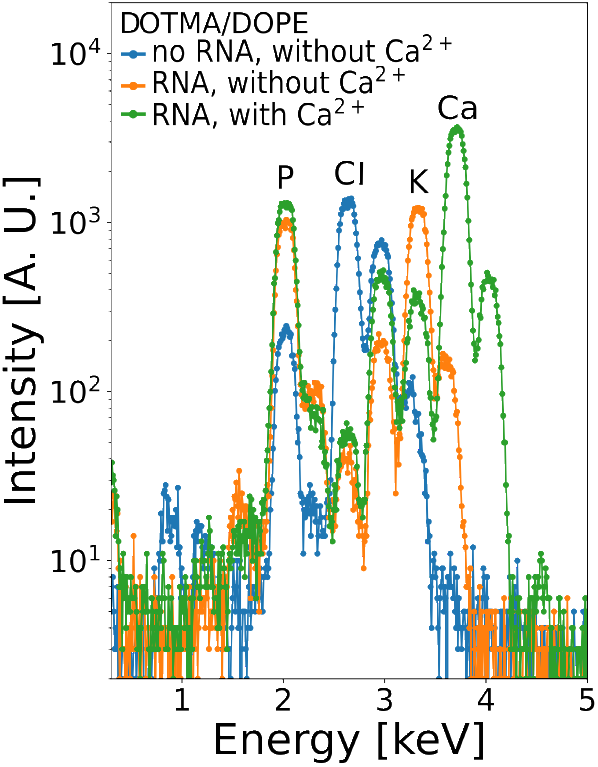
TRXF spectra of a DOTMA/DOPE monolayer without mRNA (blue), with mRNA under Ca^2+^-free conditions (orange), and with mRNA in the presence of Ca^2+^ (green). Labels indicate the dominant X-ray fluorescence emission lines of the chemical elements of interest. Equivalent figures for DOTAP/POPC and DOTAP/DOPE are presented in the Supporting Information (Fig. S4 and S5).

The P line intensity contains information on the lateral packing density of the lipid layer, due to the signal from the phospholipids in the monolayer, and of the adsorbed mRNA, which contains one phosphate group per nucleotide. For our purposes, the lateral packing density of the transfection lipid, Γ_CL_, is of interest. It is readily obtained from the packing density of the phospholipids, 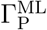, and from the known lipid mixing ratio (2:1 for DOTMA/DOPE and 1:1 for DOTAP/DOPE and DOTAP/DOPC). Table 2 provides a summary for all lipid compositions investigated. For DOTMA/DOPE, which was applied at a 2:1 molar ratio, the lateral density (Γ_CL_ = 1.05 nm^−2^) is higher than for the two other lipid systems applied as a 1:1 mixture (Γ_CL_ = 0.87 nm^−2^ for DOTAP/DOPE and Γ_CL_ = 0.76 nm^−2^ for DOTAP/DOPC). We note the higher lateral density of the DOTAP in the presence of DOPE compared to DOPC, which reflects the larger head group cross section of PC in comparison to PE and thus underlines the accuracy of the measurements.

**Table 2:**
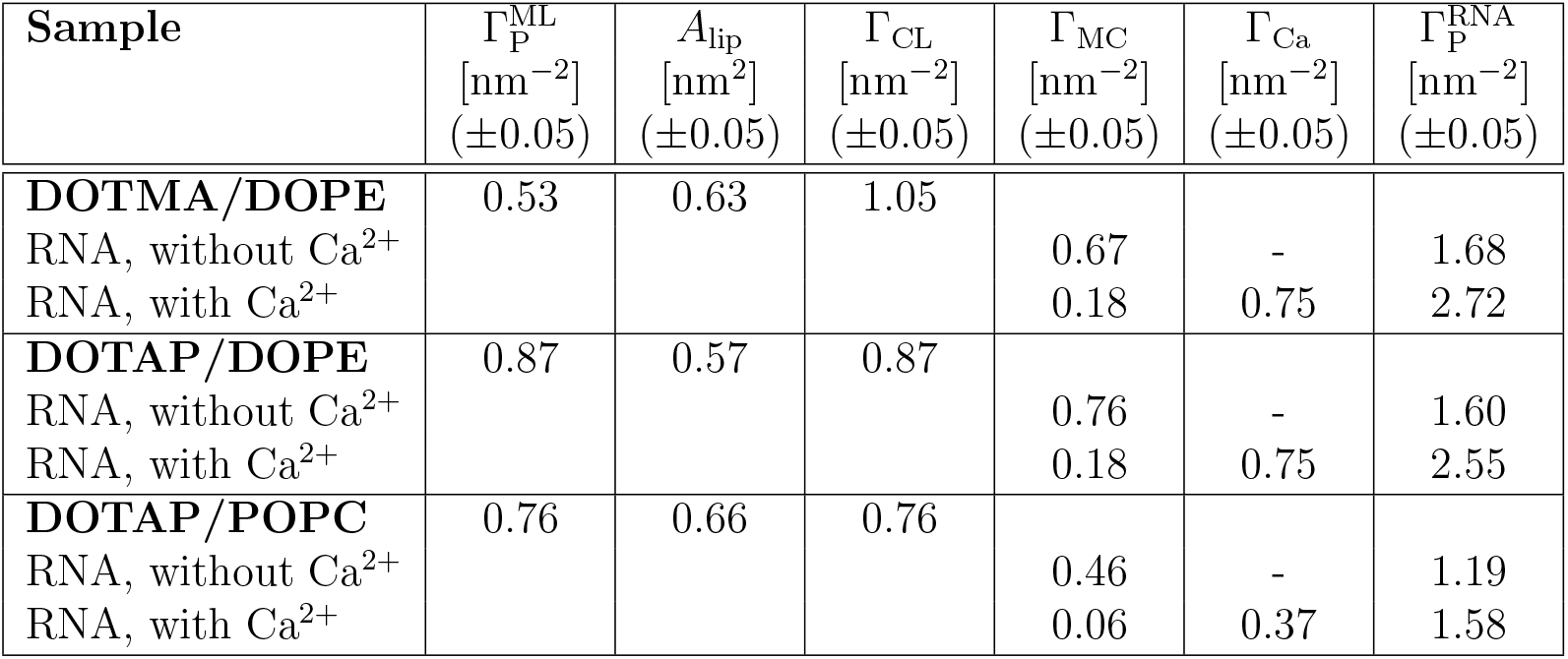
Interfacial densities of monolayer-related and mRNA-related P atoms, and of the transfection lipids, as well as the area per lipid and the surface excesses of monovalent cations (MC) and of calcium, all deduced from the TRXF spectra. See Methods section for the definition of the error estimates.

As seen in Fig. 3, adsorption of mRNA not only leads to an increase in the P line intensity but also to a dramatic decrease of the anion line intensity (here: Cl). In fact, as shown in the Supporting Information (Fig. S7), the anion line intensity drops below that in monolayer-free reference spectra, demonstrating that anions at the surface are even depleted. At the same time, the cation line intensities increase substantially, indicating the adsorption of cations along with the mRNA.

The charge balance at the interface involves contributions of all charged moieties, namely trimethylamine head groups of the transfection lipid, phosphates from the nucleotides, as well as the elemental anions and cations. The phosphate and choline/ethanolamine groups of the phospholipids do not have to be considered in this balance because the phospholipids are overall charge-neutral. For a DOTMA/DOPE monolayer, Fig. 4 shows the mutually compensating interfacial charge densities in units of elementary charges per nm^2^ (see Figs. S8 and S9 in the Supporting Information for the other lipid compositions). In the absence of mRNA, the positive charges of the lipid monolayer, whose density sets the charge density *σ*_ML_ (column on the left), are compensated by the negative charge density *σ*_A_ associated with the anions (Cl^−^ or Br^−^) from the subphase (second column from the left). In the presence of mRNA, the anions are depleted and the positive monolayer charge is instead compensated by the (excess) negative charge of the adsorbed mRNA (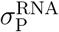, columns on the right with arrows pointing upwards), with the excess negative charge being compensated by the co-adsorbing cations (*σ*_MC_ and *σ*_Ca_, indicated with shaded boxes and arrows pointing downwards).

**Figure 4:**
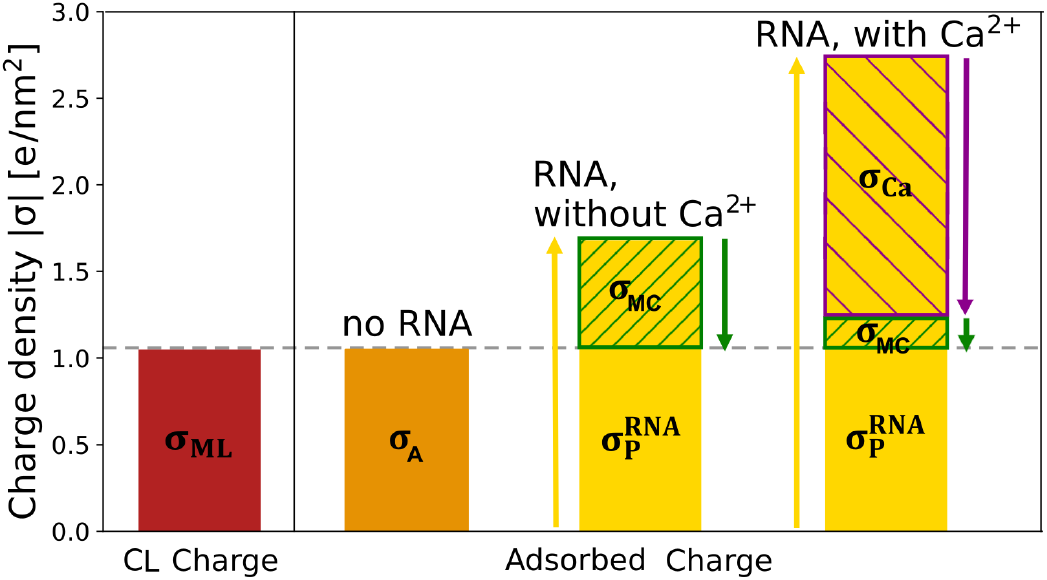
Mutually compensating interfacial charge densities related to the CL, to accumulated elemental ions, and to the adsorbed mRNA for DOTMA/DOPE in the absence and presence of RNA, with and without calcium. Data for DOTAP/POPC and DOTAP/DOPE are presented in the Supporting Information (Figs. S8 and S9).

Of particular interest is the relation between the lateral density of positively charged lipid in the monolayer, Γ_CL_, the adsorbed mRNA (encoded in 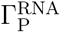), and the adsorbed ions. In LNP formulations for biological and pharmaceutical application, the N/P ratio, i.e., the ratio between nitrogen atoms of the amine groups in the (positively charged) lipids and the (negatively charged) phosphate groups from the mRNA is a key parameter. It refers to the overall molar ratio of the two oppositely charged components during manufacturing. A large excess of cationic lipid to mRNA, with N/P ratios of 4, 6 or even higher, is typically used for standard LNPs. However, no information on the local binding conditions inside the nanoparticles can be inferred from this value. In the present work we directly reveal the situation at the mRNA-lipid interface. To better reflect the conditions for the present experiments, we calculate here the stoichiometric ratio between the bound mRNA nucleotides and the CL in the monolayer, which is the inverse of the N/P ratio, and therefore denoted as P/N ratio in the following. Its values, summarized in Table 3 and seen in Fig. 5, exhibit a binding stoichiometry that strongly depends on the ionic conditions in the bulk phase and on the monolayer composition, as stated above. In contrast to the excess of CL in the overall formulation, locally we observe an excess of mRNA with respect to CL: more than one nucleotide per CL is found at the interface, with P/N ratios between 1.6 and 1.9 when only monovalent ions are present. The P/N is even higher in the presence of divalent calcium ions, between 2.1 and 2.9. The overall charge neutrality is maintained by the co-adsorption of cations together with the mRNA at the interface.

**Table 3:** Overview of the stoichiometry of charged components at the interface. See Methods section for the definition of the error estimates.

| **Sample** | $P/N = n_{\text{nucl}}/n_{\text{CL}}$<br>($\pm 0.1$) | $n_{\text{MC}}/n_{\text{CL}}$<br>($\pm 0.05$) | $n_{\text{MC}}/n_{\text{nucl}}$<br>($\pm 0.05$) | $n_{\text{Ca}}/n_{\text{CL}}$<br>($\pm 0.05$) | $n_{\text{Ca}}/n_{\text{nucl}}$<br>($\pm 0.05$) |
| --- | --- | --- | --- | --- | --- |
| **DOTMA/DOPE** |  |  |  |  |  |
| RNA, without $\text{Ca}^{2+}$ | 1.60 | 0.64 | 0.41 | - | - |
| RNA, with $\text{Ca}^{2+}$ | 2.59 | 0.17 | 0.08 | 0.71 | 0.24 |
| **DOTAP/DOPE** |  |  |  |  |  |
| RNA, without $\text{Ca}^{2+}$ | 1.84 | 0.87 | 0.47 | - | - |
| RNA, with $\text{Ca}^{2+}$ | 2.93 | 0.21 | 0.07 | 0.86 | 0.29 |
| **DOTAP/POPC** |  |  |  |  |  |
| RNA, without $\text{Ca}^{2+}$ | 1.56 | 0.60 | 0.38 | - | - |
| RNA, with $\text{Ca}^{2+}$ | 2.07 | 0.08 | 0.03 | 0.50 | 0.27 |

**Figure 5:**
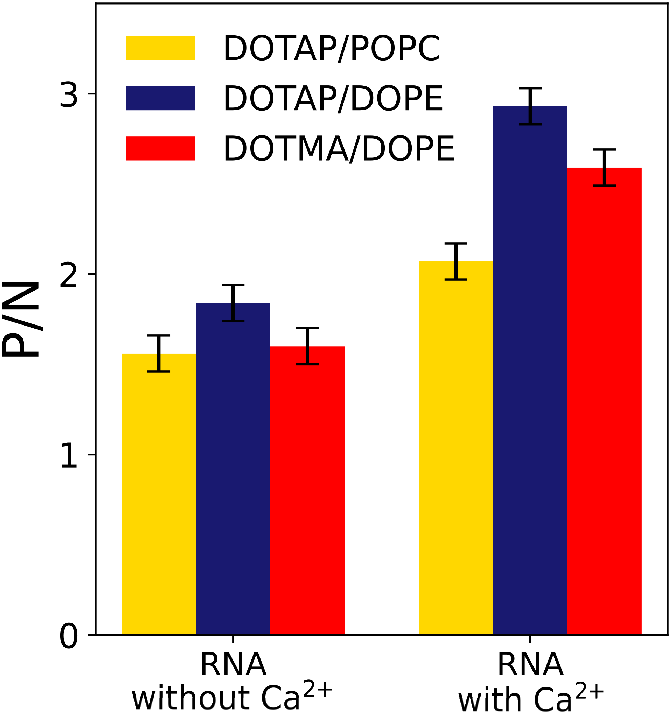
Local nitrogen-to-phosphate (N/P) ratio at the lipid/mRNA interface for all lipid compositions in the presence and absence of calcium. See Methods section for the definition of the error bars.

The amount of excess mRNA relative to the CL is very similar for all three lipid compositions investigated, but it strongly depends on the type of ions in the subphase (monovalent or divalent). The cations binding to the mRNA effectively reduce its charge density. Despite being lower in concentration, the calcium ions lead to a larger reduction of the mRNA’s charge density, so that more mRNA per cationic lipid is bound in this case. At the same time, a depletion of the monovalent cations by the divalent calcium ions is observed, demonstrating the high binding affinity of the calcium ions to the mRNA. Note that such ion binding likely occurs already to the mRNA in solution. Therefore the present measurements can give experimental information on the competing affinity of ions to the mRNA in in the bulk phase. In accordance with the requirement for overall charge neutrality, the P/N ratios are fairly well correlated with the lateral charge density in the monolayers, indicated with orange and blue straight lines in Fig. 6 A for experiments without and with calcium, respectively. For a given lipid composition, the increase of the adsorbed mRNA amount with the presence of Ca^2+^ ions (Fig. 6 B) correlates well with the thickness increase determined by GIXOS (Fig. 2 C). Here, the slope of the linear relation is characteristic for each composition (indicated with differently colored straight lines in Fig. 6 B) and systematically depends on the monolayer charge density: The higher the lateral density of CL the more compact is the RNA adsorption layer.

**Figure 6:**
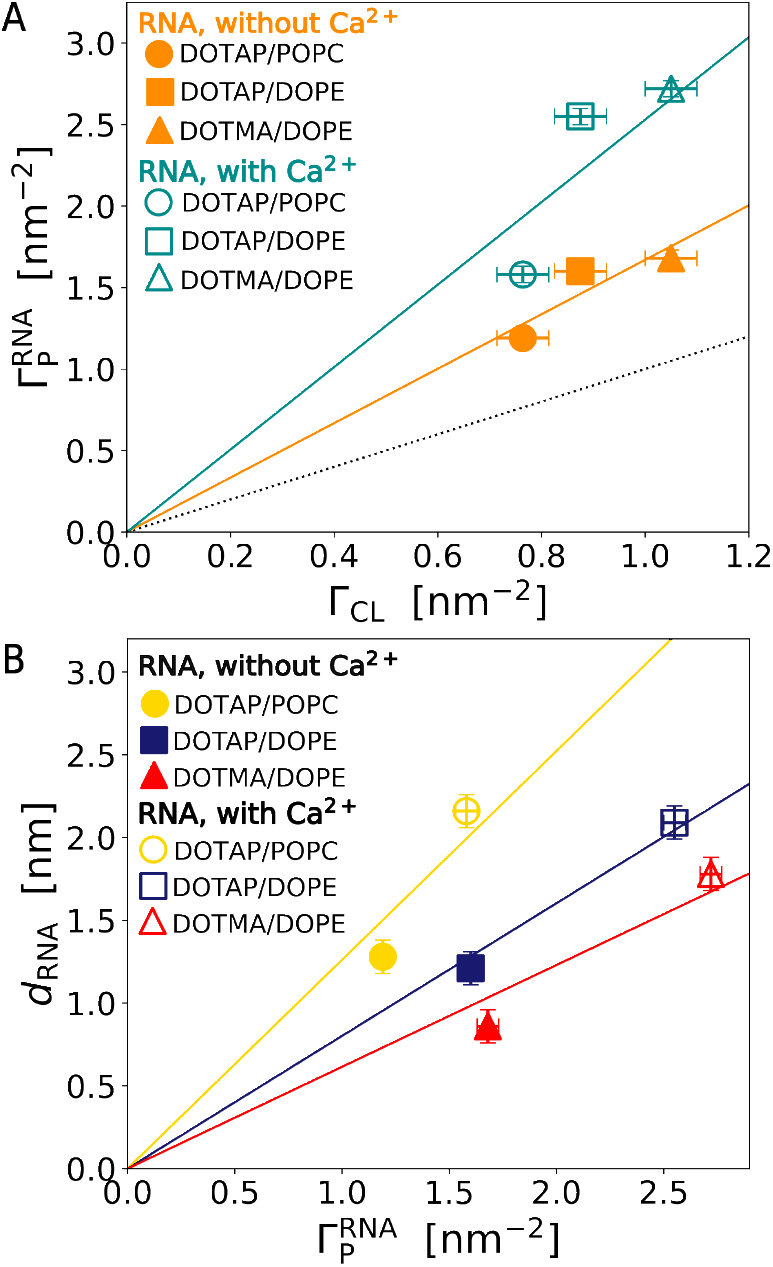
(A) 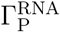 as a function of Γ_CL_. The dotted line (slope 1) indicates mRNA/CL 1:1 charge compensation. The solid lines are linear fits through zero to all data points in the absence of Ca^2+^ (slope *≈* 1.6 *⇒*60% overcompensation) and in the presence of Ca^2+^ (slope *≈* 2.5 *⇒*150% overcompensation), respectively. (B) *d*_RNA_ as a function of 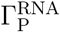. The lines are linear fits through zero to all data points for DOTAP/POPC, DOTAP/DOPE, and DOTMA/DOPE, respectively. See Methods section for the definition of the error bars.

The discrepancy between the local and the overall stoichiometry suggests that LNPs formulated at an excess of CL likely consist of two types or phases of lipidic material, of which only one is complexed to the mRNA. In fact, small-angle X-ray scattering (SAXS) measurements have pointed towards the presence of two different types of ordered material inside the LNP formulations.^13^ Such different structures may either coexist within individual LNPs or correspond to different LNP sub-populations, which are ‘mRNA-loaded’ or ‘empty’. These aspects are of high relevance for product evaluation of product quality and are currently being investigated by various groups. ^32^

### Joint GIXOS and TRXF analysis for the mRNA layer

The GIXOS and TRXF data provide complementary information, which allows to obtain an accurate and comprehensive model for the mRNA binding to the lipid layer, as schematically illustrated in Fig. 7. Specifically, we combine the information on layer thickness from GIXOS with the values for the lateral density of chemical components from TRXF, including the CL, the mRNA nucleotides, and the counterions. From the RNA layer thickness *d*_RNA_, the surface density of nucleotides (identical to 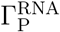), and the volume per nucleotide^33^ we calculate the layer composition in terms of the volume fractions of mRNA and water (*ϕ*_nucl_ and *ϕ*_wat_, respectively), the number of water molecules per nucleotide (*n*_wat_*/n*_nucl_), and the concentrations of monovalent and divalent cations (see Methods section for the details). The values are given in Table 4. In all cases we find a rather high packing density of the mRNA, with volume fractions varying between almost 60% and less than 30%, depending on the the monolayer composition and the ion conditions. Assuming that the rest of the volume is occupied by water molecules, at the highest mRNA packing density less than 10 water molecules are present per nucleotide. Because this number already comprises water molecules that are tightly bound to the lipid head groups, the number of free water molecules around the mRNA is even smaller. In fact, earlier bulk phase measurements on lipoplex nanoparticles are in accordance with such a low water content in the mRNA layer.^12^

**Table 4:** Overview of the chemical composition of the RNA adsorption layers at the interface in terms of nucleotide and water volume fractions (*ϕ*_nucl_ and *ϕ*_wat_, respectively), the nucleotide hydration *n*_wat_*/n*_nucl_ and the local ion concentrations *c*_MC_ and *c*_Ca_, respectively. See Methods section for the definition of the error estimates.

| Sample | $\phi_{nucl}$<br>($\pm 0.05$) | $\phi_{wat}$<br>($\pm 0.05$) | $n_{wat}/n_{nucl}$<br>($\pm 3$) | $c_{MC}$ [M]<br>($\pm 0.05$) | $c_{Ca}$ [M]<br>($\pm 0.05$) |
| --- | --- | --- | --- | --- | --- |
| **DOTMA/DOPE** |  |  |  |  |  |
| RNA, without Ca<sup>2+</sup> | 0.57 | 0.43 | 7 | 1.3 | - |
| RNA, with Ca<sup>2+</sup> | 0.46 | 0.54 | 12 | 0.17 | 0.70 |
| **DOTAP/DOPE** |  |  |  |  |  |
| RNA, without Ca<sup>2+</sup> | 0.40 | 0.60 | 15 | 1.05 | - |
| RNA, with Ca<sup>2+</sup> | 0.37 | 0.63 | 17 | 0.15 | 0.60 |
| **DOTAP/POPC** |  |  |  |  |  |
| RNA, without Ca<sup>2+</sup> | 0.28 | 0.72 | 26 | 0.60 | - |
| RNA, with Ca<sup>2+</sup> | 0.22 | 0.78 | 35 | 0.05 | 0.30 |

**Figure 7:**
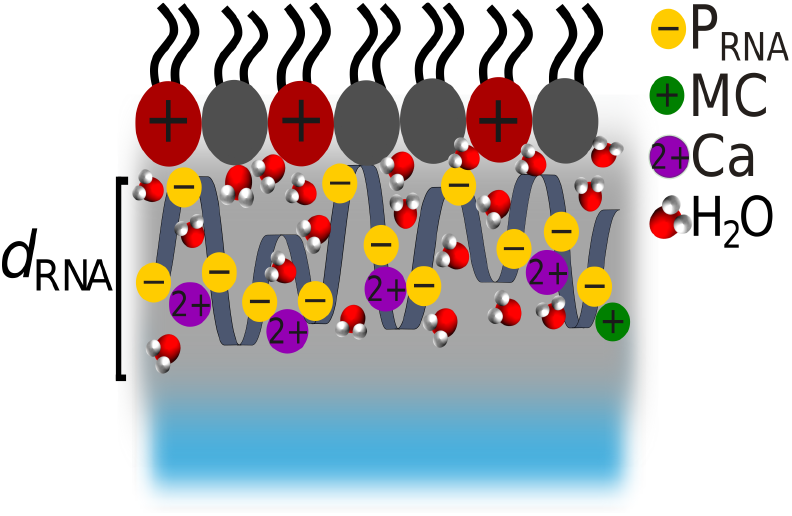
Schematic illustration of the thickness and chemical composition of the RNA adsorption layer.

As shown in Fig. 8, several important characteristics of the mRNA adsorption layer exhibit a systematic dependence on the monolayer’s positive charge density, i.e., on Γ_CL_. While the layer thickness decreases with increasing Γ_CL_ (Fig. 8 A), the mRNA packing density increases (Fig. 8 B). The highest packing density is thus obtained for mRNA binding to the DOTMA/DOPE monolayer, which has highest lateral density of the CL. Interestingly, the lipid membrane composition which results in the highest mRNA packing density also showed highest activity for spleen targeting of mRNA lipoplexes,^25^ and pharmaceutical products on this basis are currently undergoing clinical trials for cancer immunotherapy. ^28^ As the lateral density of the nucleotides increases, the number of water molecules per nucleotide decreases systematically with increasing Γ_CL_ (Fig. 8 C), from around 30 down to around 10. Also the concentrations of cations in the adsorption layer systematically increase with increasing Γ_CL_ (Fig. 8 D), as the result of the increasing lateral density of the nucleotides to which the cations bind. The concentration of monovalent cations in the mRNA layer corresponds to a concentration of up to 1.3 M (see Table 4). The divalent Ca^2+^ ions display a stronger affinity to the mRNA and largely deplete the monovalent cations from this layer, as was previously reported for the negatively charged surfaces of lipopolysaccharide layers.^34^ The highest local calcium concentration here is about 0.7 M, which represents an almost 1000-fold increase compared to the 1 mM concentration in the bulk phase. As mentioned, the numbers of monovalent and divalent cations per nucleotide are seen to be largely independent of Γ_CL_ within the experimental uncertainty (see Supporting Information, Fig. S10), indicating that it is their binding affinity to the nucleotides which is relevant for their concentration in the interfacial layer. For the salt conditions in the subphase applied here (5 mM for monovalent ions and 1 mM for calcium ions) we find a binding ratio for the monovalent cations *n*_MC_*/n*_nucl_ *≈* 0.42, and *n*_Ca_*/n*_nucl_ *≈* 0.27 with divalent calcium, see also Table 3. In addition to the charge density, also the molecular properties of the lipid monolayer are seen to influence mRNA adsorption. The two monolayers containing PE lipids adsorb more mRNA than the one containing PC lipids (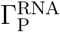 in Table 2) and the two monolayers containing DOTAP adsorb thicker mRNA layers than the one containing DOTMA (Fig. 2 C).

**Figure 8:**
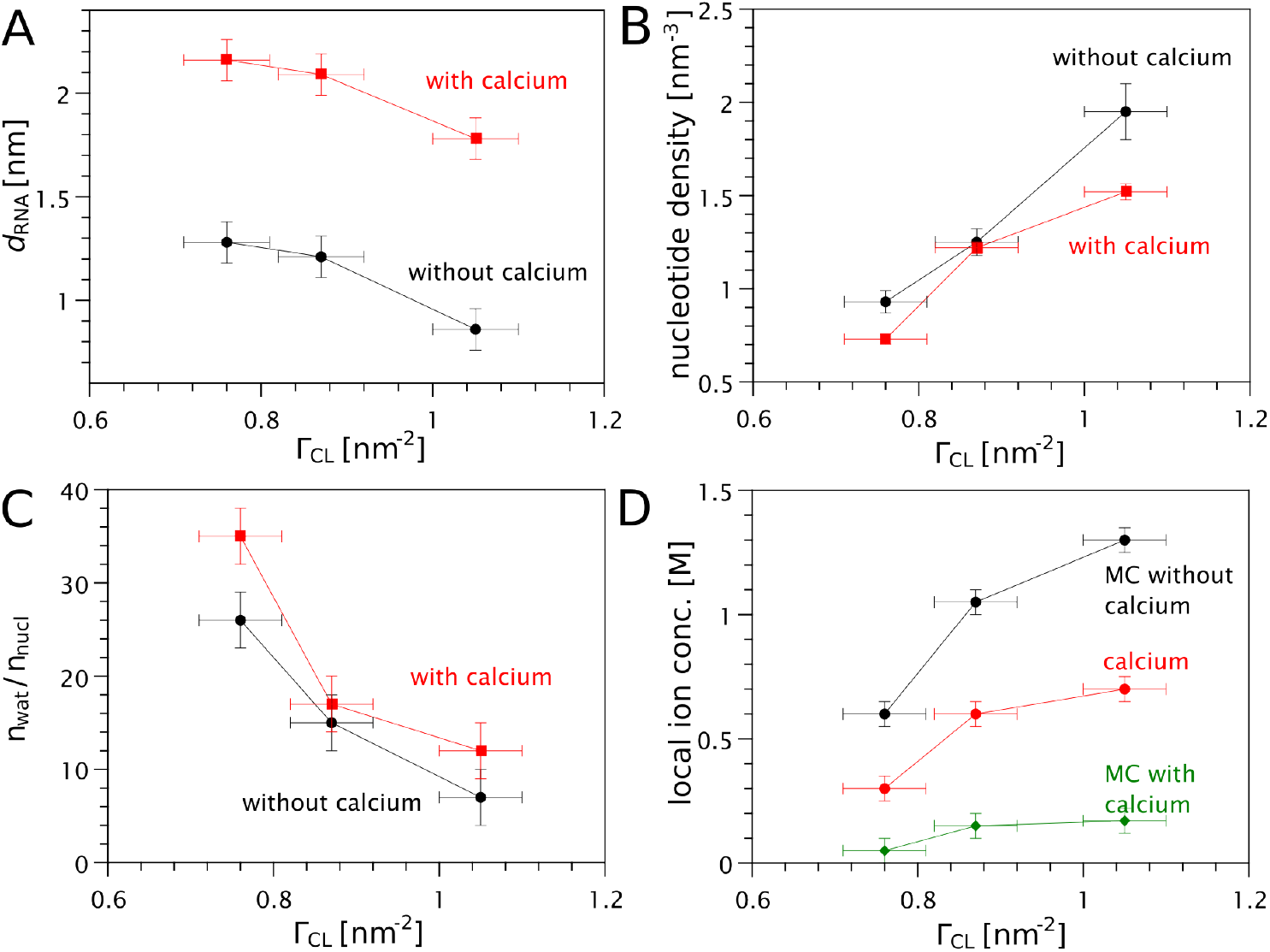
Dependence of various RNA adsorption layer characteristic on the monolayer’s positive charge density. (A) Layer thickness. (B) Nucleotide packing density. (C) Number of water molecules per nucleotide. (D) Local concentrations of cations in the adsorption layer. See Methods section for the definition of the error bars.

We note that our findings for the excess electron density from GIXOS (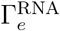, Table 1) are qualitatively in accordance with the TRXF results discussed in the previous paragraphs. With the known number of electrons for the nucleotides and water and their volume fractions from TRXF, the (excess) electron density in the mRNA layer can be calculated and compared to the experimental results. Both the relative difference between the different monolayer systems and the comparison of monovalent and divalent ions show similar trends (see Table S5 and Figs. S11 and S12 in the Supporting Information). Discrepancies must be attributed to the balance of displaced and adsorbed ions, whose electron density contribution can be substantial, and to the fact that our analysis of the GIXOS data is more sensitive to the shape of the electron density profile than to the absolute electron density plateau values, which underlines the value of combining the two methods for investigating the interfacial layers.

## Discussion

With our measurements we have elucidated the composition and structure of mRNA layers bound to oppositely charged lipid layers with unprecedented precision. The results deepen the insights and complement the results of earlier bulk phase studies of the mRNA-containing compartment in LNPs. Although the lipid monolayers constitutes only “half a bilayer”, the general findings on binding preferences can be considered to reflect to a large extend the conditions of the mRNA compartment in lipid nanoparticles. The mRNA forms a tight, discrete layer, closely attached to the lipid membrane. Its composition is driven by the balance of interactions between positive and negative charges coming from the lipids, the mRNA, and charged species from the bulk phase. In accordance with earlier bulk phase measurements, we find a defined stoichiometry between CL and mRNA, which, however, is not strictly one-to-one and does not necessarily involve direct local pairing between mRNA and cationic lipid. It is rather the overall electrostatic charge balance which drives the composition of the interfacial layer. Cations from the subphase binding to the mRNA lead to a decrease of the effective charge density and to a P/N value greater than one. Divalent cations (Ca^2+^) lead to a much higher P/N ratio than monovalent ions alone, which underlines the high binding affinity of Ca^2+^ to RNA. Also steric reasons prevent formation of commensurable lattices between mRNA and cationic lipids. As shown in the Supporting Information, the average next-neighbor distance between two CL charges in the lipid layers is 1.0 - 1.2 nm. The negative charges of the mRNA, on the other hand, are equally spaced with a distance of *≈* 0.5 nm. This complies with the substantial excess of mRNA charges with respect to the charges of the cationic lipid. We note that polyelectrolyte adsorption may in general induce some in-plane segregation of charged lipids that could lead to a more balanced interaction stoichiometry, but our observations suggest that this mechanism, which was discussed earlier,^35^ is of minor importance here. Further to the electrostatics as key driving force for the composition of the mRNA layer, here also molecular effects of the lipid layer composition were clearly discriminated, which provides valuable information for comparison of different formulations in practical development.

The conditions inside the RNA adsorption layer are decisive for central quality aspects including stability and biological activity. Chemical degradation of lipids and mRNA as well as adduct formation directly depend on the local chemical environment of the molecular moieties.^14^ It has also been shown that biological activity depends on salt conditions and on mRNA/lipid interactions.^36^

The information on the correlation between chemical characteristics of the lipid layer, the ion conditions, and mRNA binding to the lipid layers obtained here offers invaluable basic information for rational LNP design and engineering. In particular also the quantitative insights into the accumulation of ions in the mRNA layer provide a basis for evaluating manufacturing conditions and product quality. Binding stoichiometry, mRNA packing density, layer thickness, and local ionic environment can be adjusted by selecting appropriate lipid formulations and process conditions. Ions, on the one hand, can help to increase loading efficacy of the mRNA to the nanoparticles, which may influence as well endosomal processing and release. On the other hand, ions can be unfavorable for product stability, because they may accelerate chemical degradation. Regular LNP manufacturing typically involves a buffer exchange, where the particles are initially formed in a more acidic buffer than in the final product.^37^ Therefore, with the quantitative information on ionic conditions in the mRNA layer as a function of boundary conditions, certain desired parameters of the nanoparticle conformation could be tailored already during complexation such that they are maintained in the final buffer. For example, high mRNA loading may be obtained in the initial manufacturing step and undesired charged species could then be removed by subsequent buffer exchange. Due to the strong ion accumulation effect, great care should be taken to efficiently remove undesired charged species, since even low concentrations in the free buffer medium may have significant effects. Further systematic measurements will be helpful to provide deeper understanding on these quality-indicating parameters. These should include the systematic variation of ion concentration and type, pH-dependent studies including also ionizable lipids, and the investigation of the effects of different buffer molecules and chelators.

## Conclusions

Using Langmuir lipid monolayers as model systems, we accurately elucidated structure and composition of mRNA layers adsorbed to charged lipid layer surfaces. The GIXOS and TRXF measurements at the air/water interface applied here are an exquisite method to investigate the moieties binding from the aqueous subphase to the interfacial layer. Insight into RNA adsorption as a function of lateral density of the cationic transfection lipids, the chemical nature of the cationic and helper lipids and, importantly, of the ionic environment in the subphase is obtained. The competing interactions of charged moieties are the determining parameters for structure and composition of the mRNA layer attached to the lipid membrane.

The information on the mRNA packing density, the amount of water, and the accumulation of ions in the adsorbed layer is valuable for assessing stability and quality attributes of mRNA LNP formulations. Future investigation will comprise investigation of the effects of other (ionizable) lipids, the pH responsiveness and effect of buffer and chelator molecules on the composition of the mRNA layer. Such insight will provide a knowledge basis for the rational development of improved formulations and manufacturing pathways.

## Materials and methods

### Chemicals

The transfection lipids 1,2-dioleoyl-3-trimethylammonium-propane and 1,2-di-O-octadecenyl-3-trimethylammonium-propane (DOTAP and DOTMA, respectively) and the phospholipids 1-palmitoyl-2-oleoyl-glycero-3-phosphatidyl-choline and 1,2-dioleoyl-sn-glycero-3-phosphatidyl-ethanolamine (POPC and DOPE, respectively), were purchased from Sigma-Aldrich (Merck KGaA, Germany), see Fig. 1 C. Messenger RNA (mRNA) with *≈* 3900 nucleotide bases was a donation from BioNTech SE and produced by internal protocols in the form of an aqueous solution containing 10 mM HEPES and 0.1 mM EDTA. The mRNA concentration was determined by UV absorption spectrometry as *c*_RNA_ = 1.67 mg/mL. UV–vis spectra were recorded with a Lambda 650 spectrometer (PerkinElmer, Waltham, MA, USA). For the measurements, 200 *µ*L of ethanol, 500 *µ*L of mRNA solution and 200 *µ*L of 30% zwittergent were added to PMMA cuvettes with a path length of 1 cm. The Lambert-Beer law was used to determine the concentration from the attenuation at 260 nm, using the extinction coefficient of *ϵ* = 30 mL/(mg*·*cm).^38^ Ethanol, zwittergent, potassium bromide (KBr), potassium chloride (KCl), calcium bromide (CaBr_2_), calcium chloride (CaCl_2_), chloroform, methanol, and ethanol were purchased from Sigma-Aldrich (Merck KGaA, Germany) and used without further purification.

### Sample preparation

Mixtures of DOTAP and POPC (1:1 molar ratio, termed DOTAP/POPC in the following), of DOTAP and DOPE (1:1 molar ratio, termed DOTAP/DOPE in the following) and of DOTMA and DOPE (2:1 molar ratio, termed DOTMA/DOPE in the following) were prepared by first dissolving the pure components at concentrations of 1 mg/ml in chloroform and subsequent mixing of the solutions in suitable fractions. Aqueous salt solutions contained 5 mM KCl or 5 mM KBr. In some cases (termed “with calcium”) they additionally contained 1 mM CaCl_2_ or 1 mM CaBr_2_, respectively. The mRNA was dissolved in these salt solutions such that the concentration of mRNA was 0.1 mg/ml. Organic solutions of DOTAP/POPC, DOTAP/DOPE, or DOTMA/DOPE were then spread with a Hamilton syringe onto an interface between air and the above aqueous salt solutions with or without mRNA. After solvent evaporation and allowing mRNA adsorption to equilibrate for 60 min, the lipid monolayers were laterally compressed until reaching a final lateral pressure of Π = 30 mN/m, with a movable barrier at a constant compression rate of d*A*_lip_*/*d*t ≈* 0.04 nm^2^/min. To ensure adsorption equilibrium, one hour was let pass before starting the x-ray measurements. Indeed, no changes were observed when measurements were repeated at a later point.

### Grazing-incidence X-ray scattering techniques

Grazing-incidence X-ray scattering experiments (GIXOS and TRXF, see below) were carried out with the Langmuir trough GID setup ^39^ at the beamline P08 ^40^ at the storage ring PETRA III (DESY, Hamburg, Germany). The Langmuir trough (customized G4, Kibron Inc, Finland) was located in a hermetically sealed container with Kapton windows, and the temperature was kept at 20 °C by a thermostat. The container was constantly flushed with a stream of humidified helium (He) to prevent air scattering and the generation of reactive oxygen species. The synchrotron X-ray beam was monochromatized to a photon energy of 15 keV, corresponding to a wavelength of *λ* = 0.827 Å. The incident angle was adjusted to *θ*_*i*_ = 0.07 °, below the critical angle of total reflection, *θ*_*c*_ = 0.082 °. A ground glass plate was placed approximately 0.8 mm beneath the illuminated area of the monolayer in order to reduce mechanically excited surface waves. The beam footprints on the water surface were 0.25 mm x 60 mm (for GIXOS) and 1 mm x 60 mm (for TRXF), as imposed by the incident beam optics. Under total-reflection conditions an X-ray standing wave (SW) is formed at the air/water interface. The penetration depth Λ of its evanescent tail into the aqueous hemispace is a function of the angle of incidence^41^ and in the present study Λ *≃* 9 nm. The exact shape Φ(*z*) of the SW intensity along the vertical position *z* (for a given incident angle) follows from the interfacial electron density profile *ρ*(*z*), and can be calculated as described previously.^34,42^

### GIXOS

Analogous to conventional X-ray reflectometry, GIXOS allows reconstructing the interfacial electron density profile (i.e., the laterally-averaged structure of the molecular layers in the direction perpendicular to the surface) from the *Q*_*z*_-dependent scattering intensity, however at fixed incident angle. The details of this technique are described elsewhere.^19–22^ As explained more briefly in Kanduč et al.^43^ and in Grava et al., ^16^ from which the following text is partially reproduced, the *Q*_*z*_-dependence of the diffuse scattering intensity *I*(*Q*_*xy*_ ≠ 0, *Q*_*z*_) recorded at low-enough yet non-zero *Q*_*xy*_ using either an area detector (Eiger2 × 1M, Dectris AG, Switzerland) at 0.6 m distance and a beamstop inside the trough enclosure to block the totally reflected beam or with a linear detector (Pilatus 100k, Dectris AG, Switzerland, with settings described elsewhere ^24,43^). *I*(*Q*_*xy*_ ≠ 0, *Q*_*z*_) contains information equivalent to that of the conventional reflectivity *R*(*Q*_*z*_) and can be transformed as:

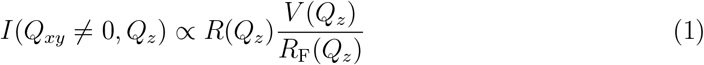

to good approximation, where *R*_F_(*Q*_*z*_) is the reflectivity of an ideal interface between the two bulk media and *V* (*Q*_*z*_) is the Vineyard function. ^44^ Note that this empirical approximation does not affect the structural analysis within the accuracy required in this study, although the capillary wave contribution to GIXOS data is not taken into account.^20^ In the present work, the GIXOS signal was measured 0.3 ° off the plane of incidence horizontally, corresponding to *Q*_*xy*_ = 0.04 Å^−1^ at *Q*_*z*_ = 0.

Like conventional reflectivity curves, theoretical GIXOS curves can be computed on the basis of an assumed interfacial electron density profile *ρ*(*z*), via the phase-correct summation of all reflected and transmitted partial waves occurring at the density gradients.^34,42^ Here, we modeled *ρ*(*z*) as a set of rough slabs representing the tail and headgroup regions of the monolayer with adjustable thickness, electron density, and roughness parameters (see Results section). In the presence of mRNA adsorbed to the monolayer surface, two additional slabs were required to reproduce the experimental GIXOS data with the model. In some cases, one additional slab for the adsorbed mRNA layer was sufficient. The electron density profile was then discretized into 1-Å-thin slices and the corresponding theoretical *Q*_*z*_-dependent reflectivities *R*(*Q*_*z*_) were calculated from the Fresnel reflection laws at each slice-slice interface using the iterative recipe of Parratt^45^ and subsequently multiplied with *V* (*Q*_*z*_)*/R*_F_(*Q*_*z*_) to obtain the theoretical GIXOS signal. For fits to the experimental data, a constant scale factor and a constant intensity background was applied to the modeled intensities, and the fitting range was limited to the consensus range of validity of Eq. 1, 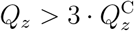, where 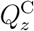 is the *Q*_*z*_-value at the critical angle of total reflection.^21^

In order to reduce the number of free fitting parameters, we tried to fix as many parameters of the HC and HG layers to their values obtained in the absence of mRNA. This approach is justified because the interaction with polynucleic acids leaves the lipid packing virtually unaffected (see Results section). Constraining all HC and HG parameters was however impossible, especially when working with aqueous solutions containing Br^−^ ions, whose high electron density affects the overall electron density profile when released upon mRNA adsorption. In the end, it was required to leave all thickness and roughness parameters free. However, *ρ*_HC_ was fixed at 300 e^−^/nm^3^ according to our previous studies^24,46^ and *ρ*_HG_ was fixed to the value obtained in the absence of mRNA. The latter constraint could not be imposed when Br^−^ was the anion, due to its more substantial electron density contribution. In view of the remaining ambiguity with regard to the exact boundary between the HG slab and the adjacent slabs describing the adsorbed mRNA layer, especially when their electron densities are similar, the most robust definition of the RNA layer thickness is: 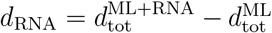, where 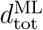 and 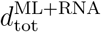 are the sums of all slab thicknesses in the absence and in the presence of mRNA, respectively.

The RNA layer’s excess electron density, 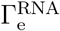, was calculated with the assumption of a constant lipid monolayer packing density. It is then simply the difference between the electron density profile of the monolayer with adsorbed mRNA, *ρ*_ML+RNA_(*z*), and the profile of the lipid monolayer without mRNA *ρ*_ML_(*z*):

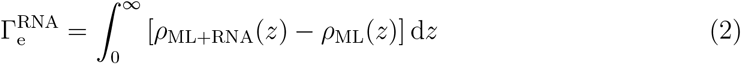

### TRXF

The x-ray fluorescence spectra were recorded using an XR-100SDD detector (Amptek, Bedford, USA). As explained in Mortara et al., ^46^ from which the following text is partially reproduced, the detector was placed almost parallel to the water surface and perpendicular to the X-ray beam axis, in order to keep elastic and Compton scattering into the detector as weak as possible. A detailed description of the method can be found elsewhere.^23,47^ The fluorescence intensity *I*_*j*_ from an element or ion species *j* is determined by the concentration profile along the direction normal to the interface *c*_*j*_(*z*):

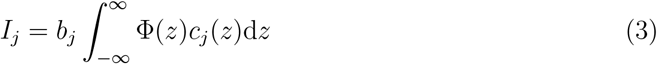

where Φ(*z*) is the SW intensity at distance *z* from the surface and *b*_*j*_ is a constant determined by the fluorescence yield and detection efficiency but independent of the structure and composition the interfacial layers.

Experimentally, *I*_*j*_ was obtained by fitting the intensity peak associated with the respective K_*α*_ emission lines in the recorded fluorescence spectra with Gaussian functions. Constraints on the peak positions were imposed based on the tabulated line energies.^48^ In cases where the K_*α*_ peak overlapped with a significant K_*β*_ peak of the neighboring element, the peaks of both elements were modeled together including the buried K_*β*_ peak, whose line energy is known and whose intensity is coupled to the respective K_*α*_ through a constant ratio that was determined independently.

The standing wave Φ(*z*) was calculated with a slab-model representation of the interfacial electron density profile,^34^ whose parameters were previously obtained in fits to the GIXOS curves. Roughness can be neglected for the computation of Φ(*z*) when *Q*_*z*_ is low, as is the case under conditions of total reflection.^23^

### Surface densities of monolayer P atoms and area per lipid

The surface densities of phosphorus (P) atoms in the lipid monolayers without adsorbed mRNA was deduced from the P fluorescence intensity 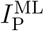 by using the justified approximation that the P atoms are located in a thin layer right in the center of the headgroup region, at *z*_P_ = *d*_HC_ + *d*_HG_*/*2. Eq. 3 then simplifies to

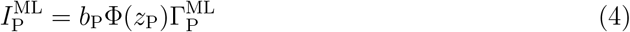

where 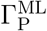 is P density in the monolayer in terms of the number of P atoms per unit area. Recalling that Φ(*z*) decays exponentially with depth *z*, this expression can be further simplified to

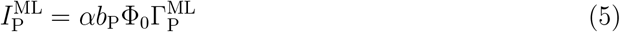

where Φ_0_ is the incident intensity and 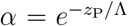 is an attenuation factor introduced for convenience. When solved for the P density, this yields

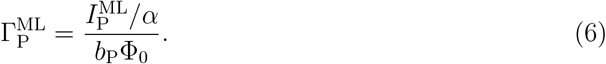

Conveniently, the experimental parameters *b*_P_ and Φ_0_ drop out when considering the ratio between two P densities:

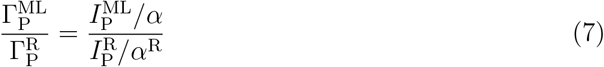

where the 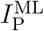 corresponds to a reference monolayer^23^ of known P density 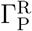, here a DSPC monolayer at Π = 30 mN/m, where 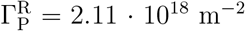 and *α*^R^ = 0.74.^49,50^ The area per lipid then follows as 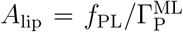, where *f*_PL_ is the phospholipid fraction (*f*_PL_ = 1*/*2 for 1:1 mixtures and *f*_PL_ = 1*/*3 for 1:2 mixtures). Similarly, the lateral density of positively charged transfection lipids is given as 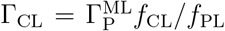, with the transfection lipid fraction *f*_CL_ = 1 − *f*_PL_. The monolayer charge density then simply follows as *σ*_ML_ = *e*Γ_CL_ (see Table 2).

### Surface densities of mRNA P atoms

The surface densities of P atoms belonging to the adsorbed mRNA were deduced from the P fluorescence intensity 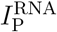 measured with the lipid monolayers in the presence of mRNA, by taking into account the known quantities 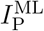 and 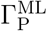 (see previous paragraph). The procedure is based on the assumption that the lateral monolayer packing, and thus 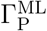, are not influenced by the mRNA adsorption (see Results section). Since P atoms from the monolayer and from the mRNA are found at different depths from the interface, where the SW intensity Φ(*z*) has different values, their contributions to 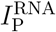 have to be modeled explicitly, as illustrated in Fig. 9. The interfacial layers are described with the electron density slab model established from the GIXOS data, with layers for the hydrocarbon chains, the headgroups, and the adsorbed mRNA. In line with our reasoning further above, in the model the monolayer P atoms are distributed over the headgroup slab of thickness *d*_HG_ at a constant concentration, see red line in Fig. 9. Similarly, the P atoms belonging to the mRNA are assumed to be evenly distributed over the slabs describing the adsorbed mRNA. Note that approximately assuming an even P distribution over the entire mRNA layer has negligible influence on the obtained results because of the comparatively long decay length of the SW.^23^ With the modeled combined concentration profile 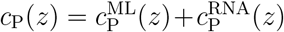 and Φ(*z*) at hand, 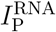 is then computed according to Eq. 3 as a function of the choice of 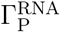. This is done on a quantitative level, since the pre-factor *b*_P_ is calibrated through the condition 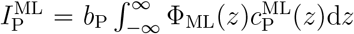, where Φ_ML_(*z*) is the SW intensity computed for the electron density profile of the monolayer without mRNA and slightly differs from that in the presence of adsorbed mRNA. In the final step, 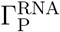 is varied until the experimental value of 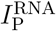 is reproduced.

**Figure 9:**
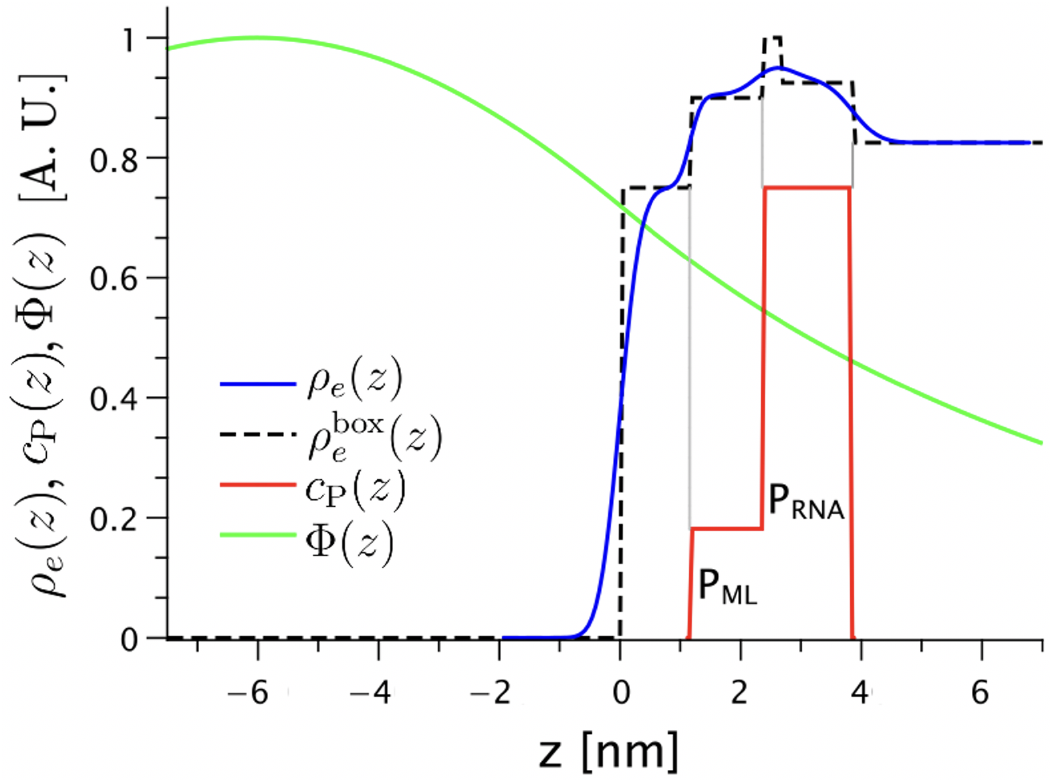
Illustration of the model used for the TRXF data analysis (here: a DOTMA/DOPE monolayer with RNA and calcium): the representation of the experimentally obtained electron density, *ρ*_*e*_(*z*), the experimentally obtained electron density without considering roughness, 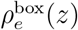, the standing wave intensity Φ(*z*), and the P concentration profile *c*_P_(*z*). All curves are normalized to have maximal values of 1, so that they can be plotted together in one graph.

### Surface excesses of anions at lipid monolayers without mRNA

According to the condition of net charge compensation, ^23^ Γ_Cl/Br_ is determined from Γ_CL_ and the constraint Γ_CL_ − Γ_Cl/Br_ = 0.

### Surface excesses of monovalent and divalent cations at lipid monolayers with adsorbed mRNA

When introducing mRNA into the salt solutions, 0.3 mM of Na^+^ are also introduced, therefore the total concentration of monovalent cations (MC) is 5.3 mM. Since we only see K^+^ from the TRXF spectra (Na^+^ is not detectable due to the low fluorescence energy), we have to apply a correction to obtain the total MC surface excess. To this end we assume that Na^+^ and K^+^ behave in the same way and that the surface excess of monovalent cations therefore is 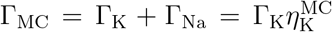, where the conversion factor between K^+^ and all monovalent cations is 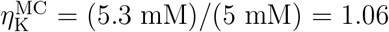. In the presence of adsorbed mRNA, we assume that co-adsorbing cations have approximately the same distribution as the P atoms of mRNA. The condition of net charge compensation was then again used to determine the cation excess and, thus, the absolute value of the cation concentration within the two RNA slabs. Namely, *b*_K_ is determined by 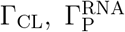, and the constraint 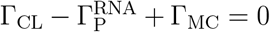. Here, we recall that the depletion of anions (Cl^−^ or Br^−^) below the bulk concentration level plays a negligible role in this balance and is therefore not considered. Finally, *b*_Ca_ is determined by 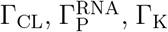, and the constraint 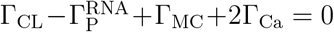.

### Volume fractions of nucleotides and water in the adsorbed RNA layer

The volume fraction of the nucleotides in the adsorption layer, *ϕ*_nucl_, is determined by the nucleotide surface density (i.e., the surface density of RNA-related P atoms), by the RNA layer thickness, and by the average nucleotide volume as 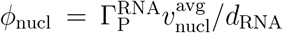, where 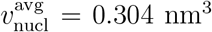 when assuming equal probabilities for all four nucleotides with volumes 0.323 nm^3^ for guanosine, 0.315 nm^3^ for adenosine, 0.291 nm^3^ for cytosine, and 0.286 nm^3^ for uridine.^33^ When neglecting the small volumetric effect of the ions, the water volume fraction simply follows as *ϕ*_wat_ = 1 − *ϕ*_nucl_.

### Number of water molecules per nucleotide and ion concentrations in the adsorption layer

The number of water molecules per nucleotide, *n*_wat_*/n*_nucl_, follows from the nucleotide surface density, from the RNA layer thickness, and from the water volume fraction as 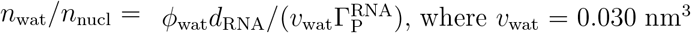 is the volume occupied per water molecule.^51^ The concentrations of monovalent and divalent cations in the RNA adsorption layer are calculated as *c*_MC_ = Γ_MC_*/d*_RNA_ and *c*_Ca_ = Γ_Ca_*/d*_RNA_, respectively.

### Error estimates

The estimated parameter uncertainties are *±* 0.1 nm for thicknesses and roughness parameters, *±* 10 e^−^nm^−3^ for electron densities, *±* 20 e^−^nm^−2^ for electron excesses, and *±* 0.05 nm^−2^ for lateral element densities. These estimates are larger than the statistical uncertainty alone, because they also account for systematic uncertainties, which are typically the dominant contribution as discussed previously.^52^ Uncertainties for other (derived) quantities were made in the spirit of Gaussian error propagation.

## Supporting Information

Pressure–area isotherms; Relative monolayer expansion before and after the injection of polynucleotides; Electron density profile parameters and plots; TRXF spectra of DOTAP/POPC and DOTAP/DOPE monolayers; Examples of full TRXF spectra; Comparison of anion x-ray fluorescence intensity of bare salt solutions and of monolayers with and without RNA; Interfacial charge balance for DOTAP/POPC and DOTAP/DOPE monolayers; Number of cations per nucleotide; Prediction of 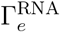 and correlation with 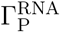; Average next-neighbor distance between two CL charges.

## Conflict of Interest

B. W. is a current employee and share holder of BioNTech SE.

H. H. is a former employee and current share holder of BioNTech SE.

## Acknowledgement

We acknowledge DESY (Hamburg, Germany), a member of the Helmholtz Association HGF, for the provision of experimental facilities. Parts of this research were carried out at PETRA III and we thank Rene Kirchhof, Milena Lippmann, and Olof Gutowski for assistance at P08 and the PETRA III chemistry lab and the use of Amptek detector, respectively. Beamtime was allocated for proposals I-20220194, I-20220952, and I-20230480. We thank Gerald Brezesinski and Lars Richter for help with the synchrotron experiments and BMBF of Germany for funding the Eiger2 × 1M detector through ErUM-Pro 05K19FK2 (Murphy, CAU Kiel).

