## supplementary text, figures, and tables for "Lipid monolayers adsorbing mRNA as models for mRNA enclosed in lipid nanoparticles for transfection"

#### – Supporting Information –

Miriam Grava,† Konrad Weber,† Joshua Reed,† Benjamin Weber,‡ Chen Shen,¶ Heinrich Haas,*,‡,§ and Emanuel Schneck*,†

†*Institute for Condensed Matter Physics, TU Darmstadt, Hochschulstraße 8, 64289 Darmstadt, Germany*

‡*BioNTech SE, An der Goldgrube 12, Mainz 55131, Germany*

¶*Deutsches Elektronen-Synchrotron DESY, Notkestrasse 85, 22607 Hamburg, Germany*

§*Department of Biopharmaceutics and Pharmaceutical Technology, Johannes Gutenberg University, 55128 Mainz, Germany*

#### Pressure–area isotherms

Fig. S1 shows pressure–area isotherms of DOTAP/DOPE monolayers in the presence of RNA with and without $Ca^{2+}$.

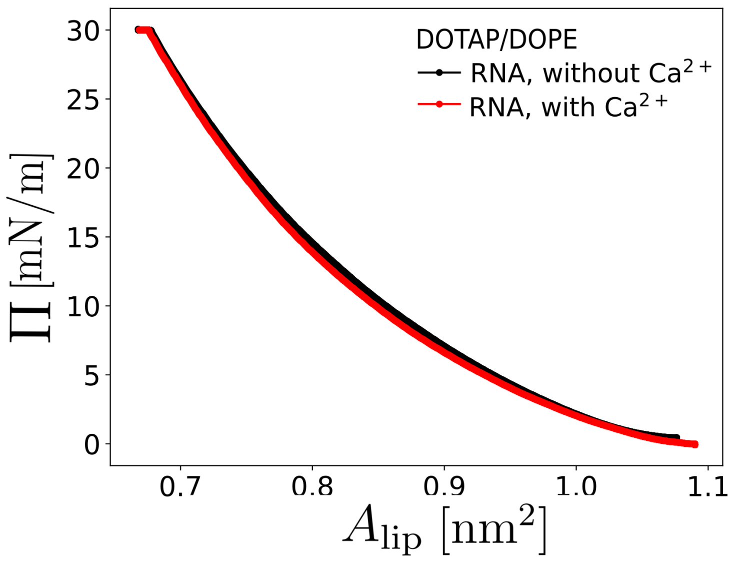


Figure S1: Langmuir isotherms of DOTAP/DOPE monolayers in the presence of RNA without $Ca^{2+}$ and with $Ca^{2+}$.

#### Relative monolayer expansion before and after the injection of polynucleotides

Table S1 summarizes the relative expansions of various monolayers in response to the injection of poly(uridylic acid) into the subphase (0.2 mg/mL final concentration). The measurements were done on 5 mM KBr subphases with a monolayer of a 1:1 mixture of di-octadecyl-TAP (DOTAP18:0) and DSPC, a monolayer of pure di-octadecyl-TAP, and a monolayer of pure DOTAP.

Table S1: Relative expansions of various monolayers in response to the injection of poly(uridylic acid).

| **Sample** | Relative expansion |
|---|---|
| DOTAP18:0/DSPC 1:1 | 100.3 % |
| DOTAP18:0 | 104.0 % |
| DOTAP | 101.6 % |

### Electron density profile parameters and plots

Tables S2, S3, and S4 summarize the slab model parameters obtained by GIXOS for DOTMA/DOPE, DOTAP/POPC, and DOTAP/DOPE, respectively, in the absence and presence of RNA, with and without calcium. Figs. S2 and S3 show the associated electron density profiles for DOTAP/POPC and DOTAP/DOPE, respectively.

Table S2: Slab model parameters obtained by GIXOS for DOTMA/DOPE. The anion was $Cl^-$. [a]Fixed at literature value. [b]Constrained to match value in the absence of mRNA. [c]One RNA slab was sufficient to model the data.

| **parameter** | no RNA without $Ca^{2+}$ | RNA without $Ca^{2+}$ | RNA with $Ca^{2+}$ |
|---|---|---|---|
| $\rho$ [$e^-$/nm$^3$] ($\pm$ 10) | | | |
| HC | 300[a] | 300[a] | 300[a] |
| HG | 360 | 360[b] | 360[b] |
| RNA, in | - | 380[c] | 400 |
| RNA, out | - | (-)[c] | 370 |
| $d$ [nm] ($\pm$ 0.1) | | | |
| HC | 1.3 | 1.2 | 1.2 |
| HG | 0.8 | 0.7 | 1.2 |
| RNA, in | - | 1.0[c] | 0.3 |
| RNA, out | - | (-)[c] | 1.2 |
| $\varepsilon$ [nm] ($\pm$ 0.1) | | | |
| air/HC | 0.2 | 0.3 | 0.3 |
| HC/HG | 0.2 | 0.1 | 0.2 |
| HG/wat | 0.1 | - | - |
| HG/RNA | - | 0.1 | 0.1 |
| RNA in/out | - | (-)[c] | 0.3 |
| RNA/wat | - | 0.5 | 0.4 |

Table S3: Slab model parameters obtained by GIXOS for DOTAP/POPC. The anion was $Br^-$. [a]Fixed at literature value.

| **parameter** | no RNA without $Ca^{2+}$ | RNA without $Ca^{2+}$ | no RNA with $Ca^{2+}$ | RNA with $Ca^{2+}$ |
|---|---|---|---|---|
| $\rho$ [$e^-$/nm$^3$] ($\pm$ 10) | | | | |
| HC | 300[a] | 300[a] | 300[a] | 300[a] |
| HG | 390 | 380 | 400 | 390 |
| RNA, in | - | 340 | - | 290 |
| RNA, out | - | 360 | - | 350 |
| $d$ [nm] ($\pm$ 0.1) | | | | |
| HC | 1.3 | 1.3 | 1.3 | 1.4 |
| HG | 0.8 | 0.8 | 0.8 | 0.6 |
| RNA, in | - | 0.2 | - | 0.6 |
| RNA, out | - | 1.1 | - | 1.7 |
| $\varepsilon$ [nm] ($\pm$ 0.1) | | | | |
| air/HC | 0.2 | 0.3 | 0.2 | 0.3 |
| HC/HG | 0.2 | 0.2 | 0.2 | 0.1 |
| HG/wat | 0.2 | - | 0.2 | - |
| HG/RNA | - | 0.2 | - | 0.5 |
| RNA in/out | - | 0.2 | - | 0.2 |
| RNA/wat | - | 0.4 | - | 0.3 |

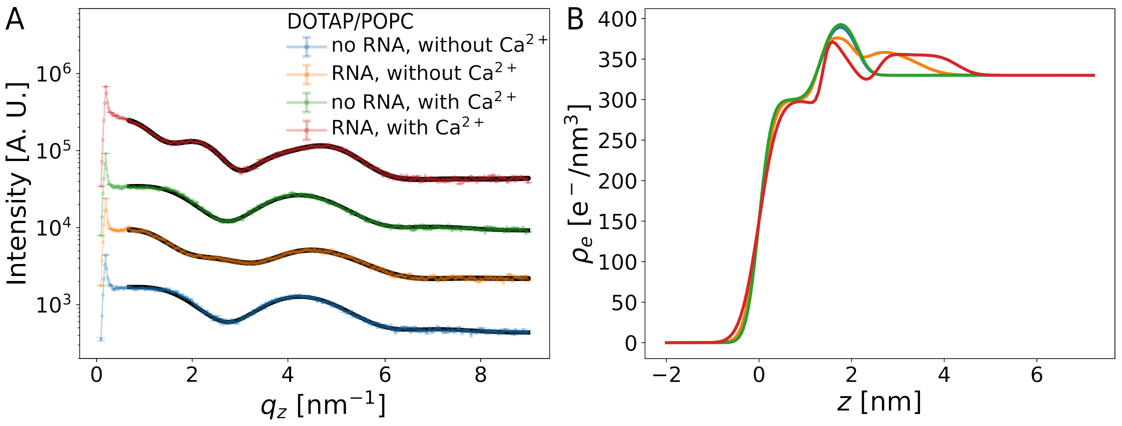


Figure S2: (A) Experimental GIXOS curves and (B) corresponding electron density profiles $\rho(z)$ for a DOTAP/POPC monolayer on different subphases. For clarity, the curves in panel (A) are vertically offset in the semi-logarithmic plot through multiplication with suitable factors (5, 20 and 100).

Table S4: Slab model parameters obtained by GIXOS for DOTAP/DOPE. The anion was $Cl^-$. [a]Fixed at literature value. [b]Constrained to match value in the absence of mRNA.

| **parameter** | no RNA without $Ca^{2+}$ | RNA without $Ca^{2+}$ | RNA with $Ca^{2+}$ |
|---|---|---|---|
| $\rho$ [$e^-/nm^3$] ($\pm$ 10) | | | |
| HC | 300[a] | 300[a] | 300[a] |
| HG | 380 | 380[b] | 400 |
| RNA, in | - | 360 | 390 |
| RNA, out | - | 370 | 370 |
| $d$ [nm] ($\pm$ 0.1) | | | |
| HC | 1.3 | 1.3 | 1.3 |
| HG | 0.8 | 0.7 | 0.7 |
| RNA, in | - | 0.3 | 0.5 |
| RNA, out | - | 1.0 | 1.7 |
| $\varepsilon$ [nm] ($\pm$ 0.1) | | | |
| air/HC | 0.2 | 0.2 | 0.3 |
| HC/HG | 0.2 | 0.2 | 0.3 |
| HG/wat | 0.2 | - | - |
| HG/RNA | - | 0.1 | 0.2 |
| RNA in/out | - | 0.3 | 0.3 |
| RNA/wat | - | 0.5 | 0.5 |

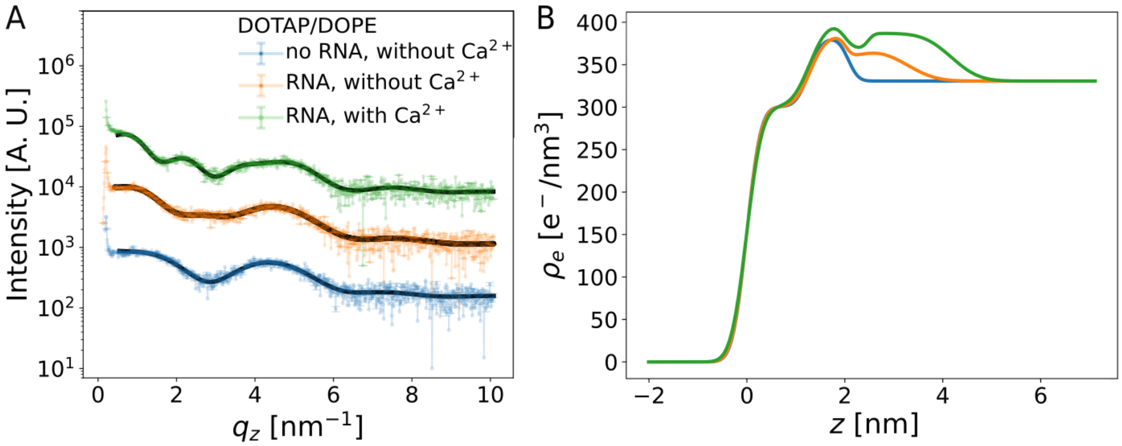


Figure S3: (A) Experimental GIXOS curves and (B) corresponding electron density profiles $\rho(z)$ for a DOTAP/DOPE monolayer on different subphases. For clarity, the curves in panel (A) are vertically offset in the plot through multiplication with suitable factors (1, 10 and 50).

### TRXF spectra of DOTAP/POPC and DOTAP/DOPE monolayers

Figs. S4 and S5 show the TRXF spectra obtained with DOTAP/POPC and DOTAP/DOPE monolayers, respectively, under various conditions.

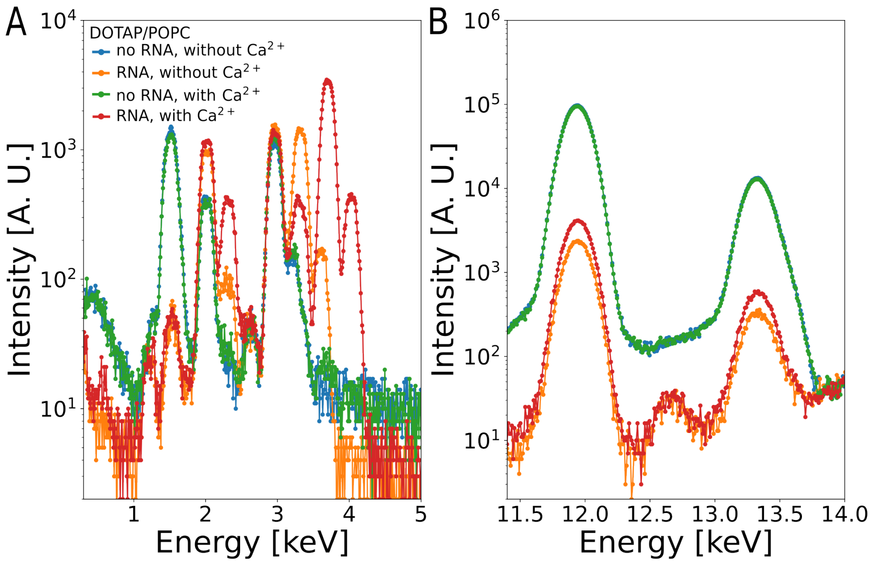


Figure S4: TRXF spectra of a DOTAP/POPC monolayer under various conditions (see legend). (A) Low energy region. (B) High energy region.

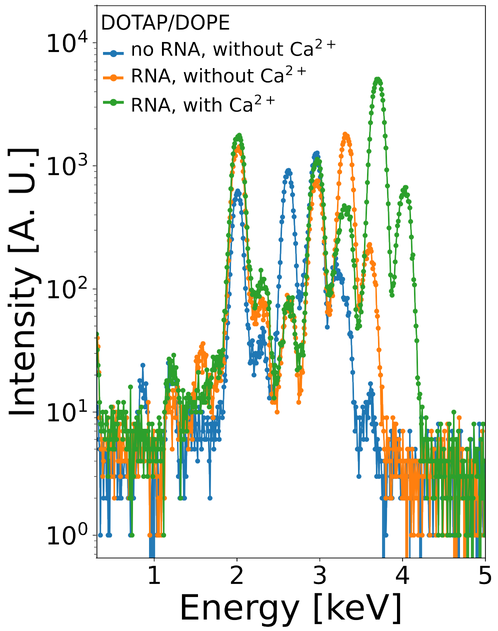


Figure S5: TRXF spectra of a DOTAP/DOPE monolayer under various conditions (see legend).

### Examples of full TRXF spectra

Fig. S6 shows a representative set of fluorescence spectra along the full energy range.

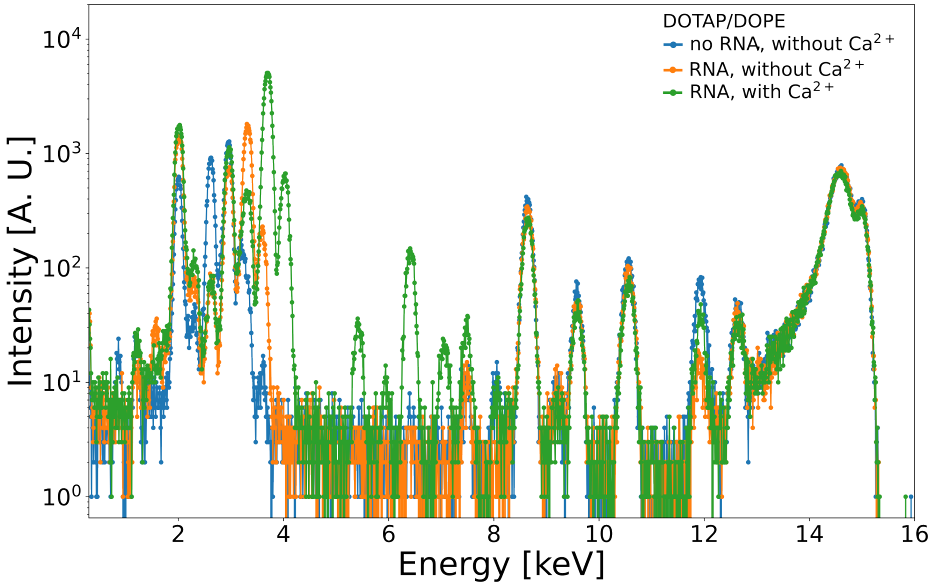


Figure S6: Full fluorescence spectrum induced via photoelectric ionization of DOTAP/DOPE lipid monolayer at $\Pi = 30$ mN/m under various conditions (see legend).

### Comparison of anion x-ray fluorescence intensity of bare salt solutions and of monolayers with and without RNA.

Fig. S7 shows the TRXF spectra obtained with a DOTAP/DOPE monolayer in the absence and presence of RNA in comparison with TRXF spectra from the surface of a bare aqueous solution of 5 mM KCl. In the presence of RNA, the chloride emission line at around 2.6 keV is weaker than that observed with the bare aqueous solution.

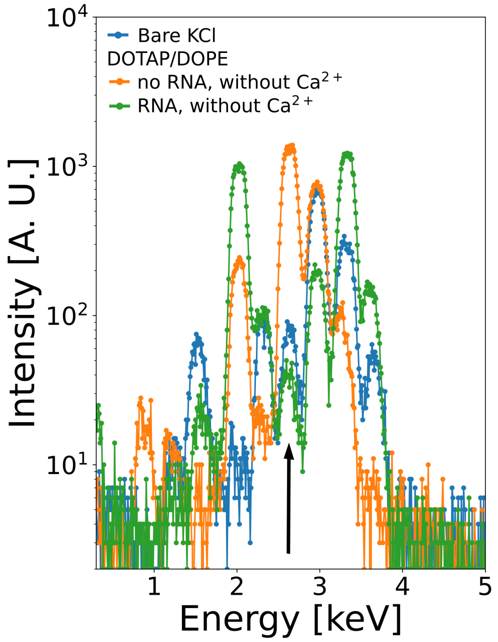


Figure S7: TRXF spectra of a DOTAP/DOPE monolayer in the absence and presence of RNA and comparison with TRXF spectra from the surface of a bare aqueous solution of 5 mM KCl. The chloride emission line at around 2.6 keV is indicated with an arrow.

### Interfacial charge balance for DOTAP/POPC and DOTAP/DOPE monolayers

Figs. S8 and S9 show the mutually compensating interfacial charge densities for DOTAP/POPC and DOTAP/DOPE monolayers, respectively, under various conditions.

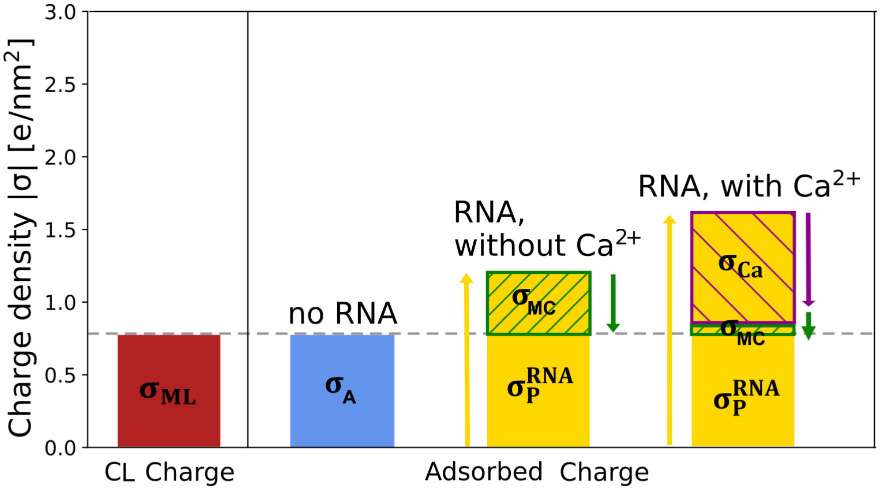


Figure S8: Mutually compensating interfacial charge densities related to the CL, to accumulated elemental ions, and to the adsorbed mRNA for DOTAP/POPC without RNA (A), with RNA without $Ca^{2+}$ (B), and with RNA with $Ca^{2+}$ (C).

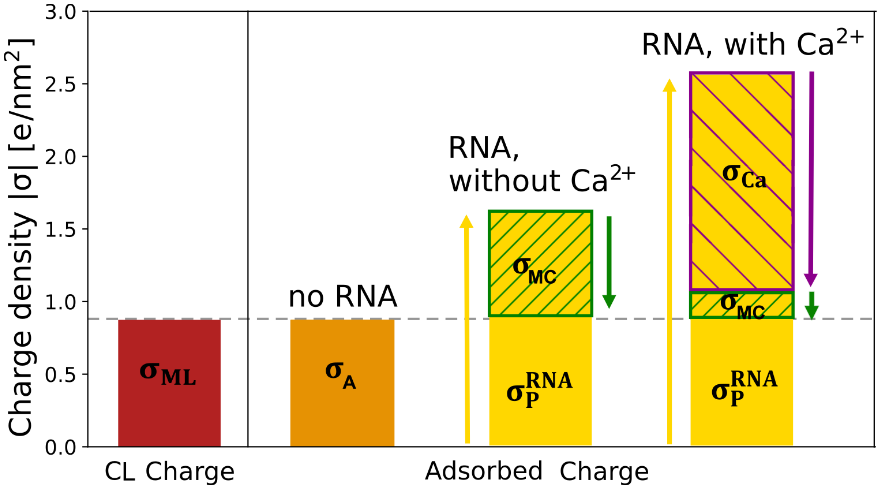


Figure S9: Mutually compensating interfacial charge densities related to the CL, to accumulated elemental ions, and to the adsorbed mRNA for DOTAP/DOPE without RNA (A), with RNA without $Ca^{2+}$ (B), and with RNA with $Ca^{2+}$ (C).

### Number of cations per nucleotide

Fig. S10 shows the numbers of monovalent and divalent cations per nucleotide as a function of the monolayer charge density $\Gamma_{CL}$ under various salt conditions.

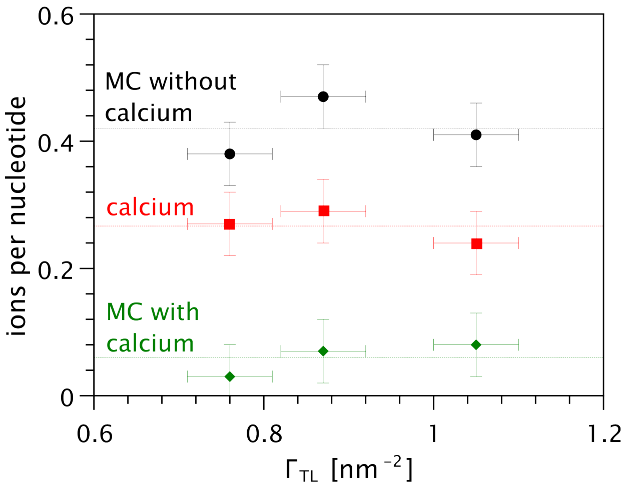


Figure S10: Numbers of monovalent cations (MC) and divalent cations per nucleotide as a function of the monolayer charge density $\Gamma_{CL}$ under various salt conditions. Horizontal lines indicate the average values.

### Prediction of $\Gamma_e^{\mathbf{RNA}}$ and correlation with $\Gamma_{\mathbf{P}}^{\mathbf{RNA}}$

The prediction of $\Gamma_e^{\mathrm{RNA}}$ starts with a calculation of the electron density of RNA. For this purpose, we count the electrons of all four nucleotides in their bound configuration (accounting for the loss of one $H_2O$ upon polymerization).

Adenosine monophosphate - $H_2O$: 171 electrons

Cytidine monophosphate - $H_2O$: 159 electrons

Guanosine monophosphate - $H_2O$: 179 electrons

Uridine monophosphate - $H_2O$: 159 electrons

On average (which we also assume for the volume), we obtain 167 electrons per nucleotide. Divided by the volume of 0.304 $\mathrm{nm}^3$ this results in an electron density of 549 $\mathrm{e}^-/\mathrm{nm}^3$ for RNA. The average electron density of the RNA adsorption layer then follows from the electron densities of RNA and of water and from their respective volume fractions $\phi_{\mathrm{nucl}}$ and $\phi_{\mathrm{wat}}$, respectively. The electron excess $\Gamma_e^{\mathrm{RNA}}$ is then $d_{\mathrm{RNA}}$ multiplied with the difference between the layer's electron density and that of pure water. The resulting predictions are summarized in Table S5 and compared to the values determined by GIXOS. The correlation between predicted and measured values is also shown in Fig. S11. By construction, a similar correlation is also found between $\Gamma_e^{RNA}$ and $\Gamma_P^{RNA}$, see Fig. S12.

Table S5: Electron excess as deduced from the GIXOS data and as predicted from the layers' chemical composition. [a]Not quantified when $Br^-$ is the anion (see main text).

| **DOTMA/DOPE** | without $Ca^{2+}$ | with $Ca^{2+}$ |
|---|---|---|
| $\Gamma_e^{RNA}$ [$e^-/nm^2$] ($\pm$ 20) | 53 | 87 |
| predicted | 111 | 179 |
| **DOTAP/DOPE** | without $Ca^{2+}$ | with $Ca^{2+}$ |
| $\Gamma_e^{RNA}$ [$e^-/nm^2$] ($\pm$ 20) | 40 | 122 |
| predicted | 105 | 168 |
| **DOTAP/POPC** | without $Ca^{2+}$ | with $Ca^{2+}$ |
| $\Gamma_e^{RNA}$ [$e^-/nm^2$] ($\pm$ 20) | (-)[a] | (-)[a] |
| predicted | 79 | 104 |

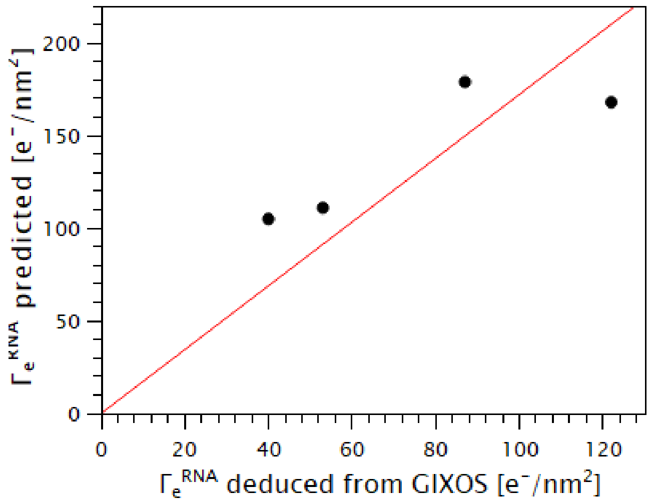


Figure S11: Correlation between predicted and measured values of $\Gamma_e^{RNA}$.

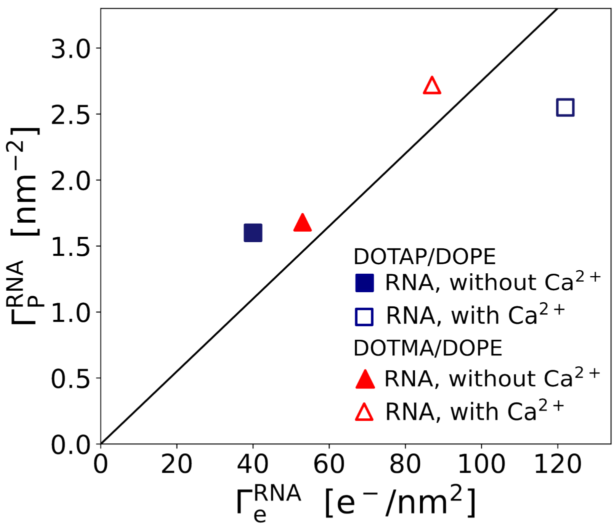


Figure S12: Correlation between $\Gamma_e^{\mathrm{RNA}}$ and $\Gamma_{\mathrm{P}}^{\mathrm{RNA}}$.

### Average next-neighbor distance between two CL charges

If, as the simplest approximation, the CL are assumed to be evenly distributed in the monolayer, then the average next-neighbor distance between two CL charges is $a = \gamma\sqrt{A_{\mathrm{lip}}/f_{\mathrm{CL}}}$, where $\gamma = 1$ for a square lattice and $\gamma \approx 1.07$ for a hexagonal lattice. With the $A_{\mathrm{lip}}$ values in Table 2 of the main text, we obtain $a = 1.0$ - $1.2$ nm, irrespective of the assumed lattice.
